# Widespread sympatry in a species-rich clade of marine fishes (Carangoidei)

**DOI:** 10.1101/2022.09.26.509594

**Authors:** Jessica R. Glass, Richard C. Harrington, Peter F. Cowman, Brant C. Faircloth, Thomas J. Near

## Abstract

The patterns of speciation in marine fishes are largely unknown, in part due to the deficiency of species-level phylogenies and information on species’ distributions, and partly due to conflicting relationships between species’ dispersal, range size, and patterns of co-occurrence. Most research on global patterns of marine fish speciation has focused on coral reef or pelagic species. Carangoidei is a clade of marine fishes including the trevallies, remoras, and dolphinfishes that utilize both coral reef and pelagic environments, spanning the ecologies of coral reef obligate and open-ocean species. We used sequence capture of 1314 ultraconserved elements (UCEs) from 154 taxa to generate a phylogeny of Carangoidei and its parent clade, Carangiformes. Age-range correlation analyses of the geographic distributions and divergence times of sister species pairs reveal widespread sympatry, with 73% of sister species pairs exhibiting a sympatric geographic distribution, regardless of node age, and most species pairs co-existing across large portions of their ranges. We also observe greater disparity in body size and water column depth utilization between sympatric than allopatric sister species. These and other ecological or behavioral attributes likely facilitate sympatry among the most closely related carangoid species, which exhibit sympatry at a larger taxonomic scale than has previously been described in marine fishes.

## Introduction

For decades, biologists have debated whether there is a universal paradigm to explain patterns and processes of speciation in marine habitats (Jordan 1905; Dobzhansky 1937; Mayr 1942; Palumbi 1994; Butlin et al. 2008; Rocha and Bowen 2008). From describing modes of speciation and mechanisms of dispersal (Butlin et al. 2008; Luiz et al. 2012), to characterizing latitudinal and longitudinal diversity gradients (Cowman 2014; Hodge et al. 2014; Siqueira et al. 2016) and hypothesizing geographic origins of diversity (Briggs 2003; Connolly et al. 2003; Cowman and Bellwood 2013), the rise of genetic methods and oceanographic modeling has upended traditional assumptions that vicariance leading to allopatry (Mayr 1942) is the default mechanism of speciation in the ocean (Rocha and Bowen 2008; Bowen et al. 2013). Although biogeographic barriers have been shown to result in allopatric speciation in certain circumstances (Schmidt 2007; Lessios 2008; Rocha and Bowen 2008; DiBattista et al. 2013; Hodge and Bellwood 2016), open-ocean environments have few obvious geographic barriers to dispersal, and other factors such as body size, ecological tolerance, pelagic larval duration, and dispersal ability may be the prominent facilitators of speciation (Palumbi 1994; Norris 2000; Luiz et al. 2012, 2015; Mora et al. 2012).

To assess contemporary and historical patterns of marine speciation and biogeography, scientists have employed a variety of approaches (Knowlton 2003; Losos and Glor 2003; Rocha and Bowen 2008; Landis et al. 2013; Matzke 2013; Hodge and Bellwood 2016). One comparative method, age-range correlation, analyzes the extent of range overlap between sister species pairs compared to the age of the phylogenetic node age immediately subtending them as a proxy of species’ age (Chesser and Zink 1994; Barraclough and Vogler 2000; Fitzpatrick and Turelli 2006; Quenouille et al. 2011; Hodge and Bellwood 2015). Age-range correlations can be examined across many sister species pairs to look for associations between geographic patterns and relative node ages. Assuming a primarily allopatric speciation model, random, independent changes in ranges over time should lead to greater sympatry at older nodes, whereas a sympatric speciation model should reflect greater range overlap in recently diverged sister species compared to more distantly related sister clades (Fitzpatrick and Turelli 2006; Hodge and Bellwood 2015). Peripatric speciation, caused when a population becomes isolated at the periphery of its ancestral distribution, can be assessed by examining range size evenness (range symmetry) between sister species and is suggested when the ranges of recently diverged sister species are highly asymmetrical due to one species having a smaller range on the edge of the larger ancestral range (Barraclough and Vogler 2000; Hodge et al. 2012).

Few studies have applied analyses of age-range correlation and range symmetry in large and taxonomically inclusive lineages of vertebrates, particularly marine fishes (Fitzpatrick and Turelli 2006; Phillimore et al. 2008; Wollenberg et al. 2011; Mora et al. 2012; Anacker and Strauss 2014; Hodge and Bellwood 2015; Tavera and Wainwright 2019). The use of such approaches is hindered by lack of comprehensive taxon sampling and availability of range data for many species. Existing age-range correlation studies on marine fishes have focused on those occupying tropical coral reefs because they are generally habitat-restricted, have relatively small body-sizes, and are a key component of the hyperdiverse Indo-Australian Archipelago (Rocha et al. 2005; Rocha and Bowen 2008; Bowen et al. 2013; Cowman 2014; Hodge and Bellwood 2016). While these methods have challenges, such as distinguishing between sympatric speciation and allopatric speciation with subsequent range changes, i.e., secondary sympatry (Chesser and Zink 1994; Barraclough and Vogler 2000; Losos and Glor 2003; Pontarp et al. 2015), broadly examining relationships between species ranges and node ages across large and diverse clades is nevertheless useful for understanding contemporary and historic biogeography in marine fishes, particularly pelagic and non-reef obligate species with high dispersal abilities. For these species, the traditional models of allopatric and parapatric speciation that are believed to affect coral reef species (Rocha and Bowen 2008; Hodge and Bellwood 2016) may be less important than processes such as colonization ability and ecological divergence through habitat partitioning or reproductive timing (Palumbi 1994; Norris 2000; Rocha et al. 2005). Examining clade-level patterns of species ranges and integrating approaches such as age-range correlation allow one to start quantifying the relationship between biogeography and speciation at a larger taxonomic scale.

In this study, we characterize contemporary patterns of biogeography in species belonging to a large clade of coastal-pelagic percomorph marine fishes, Carangoidei (Girard et al. 2020), which contains *Coryphaena* (dolphinfishes), Echeneidae (remoras), *Rachycentron canadum* (Cobia), and Carangidae (jacks and pompanos). These fishes have habitat preferences ranging from reef-associated to pelagic-neritic to brackish, although the group can broadly be classified as coastal-pelagic. The variety of life history characteristics of the carangoids frequently exclude them from studies on coral reef obligate fishes, as well as studies that focus on open-water pelagic species such as billfishes and large tunas, because they do not exhibit ecological traits characteristic of entirely one group. For example, some genera within Carangoidei (e.g., *Seriola*, *Caranx, Remora, Coryphaena*) have high dispersal potential due to their large body size, schooling activity, and association with drifting seaweed rafts, similar to pelagic fishes (Luiz et al. 2015). Yet many carangoid species also display restricted home ranges, a trait more characteristic of reef fishes (Gillanders et al. 2001; Ross and Lancaster 2002; Meyer et al. 2007; Afonso et al. 2009; Brown et al. 2010; Fontes et al. 2014). All carangoids are assumed to have pelagic larval dispersal, but length of larval drifting and juvenile settlement patterns vary across species (Smith-Vaniz 1984; Sudekum et al. 1991; Leis et al. 2006, 2007). Moreover, the importance of pelagic larval duration on dispersal, range size, and speciation rate for marine fishes remains disputed (Lester and Ruttenberg 2005; Lester et al. 2007; Weersing and Toonen 2009; Mora et al. 2012). Ecological characteristics that would favor a certain speciation mechanism (e.g., body size, migration, reef or structural associations) are either unknown or inconsistently assessed across carangoid species. For example, habitat shifts from reef to non-reef environments influence rates of carangoid morphological diversification but not rates of lineage diversification (Frédérich et al. 2016).

The regions of highest species richness of Carangoidei are the reef-abundant Indo-Australian Archipelago and the Western Indian Ocean (Fig. 1A), making carangoids important candidates for discussions on origins and historical patterns of tropical and sub- tropical fish biodiversity (Cowman 2014). Furthermore, many carangoid species are active predators that play ecologically significant roles in coral reef and coastal ecosystems (Sudekum et al. 1991; Andaloro and Pipitone 1997; Glass et al. 2020). Carangoids are thus an important group for studying biogeographic patterns of fishes that span coral reefs and coastal habitats to the open ocean.

**Figure 1.**
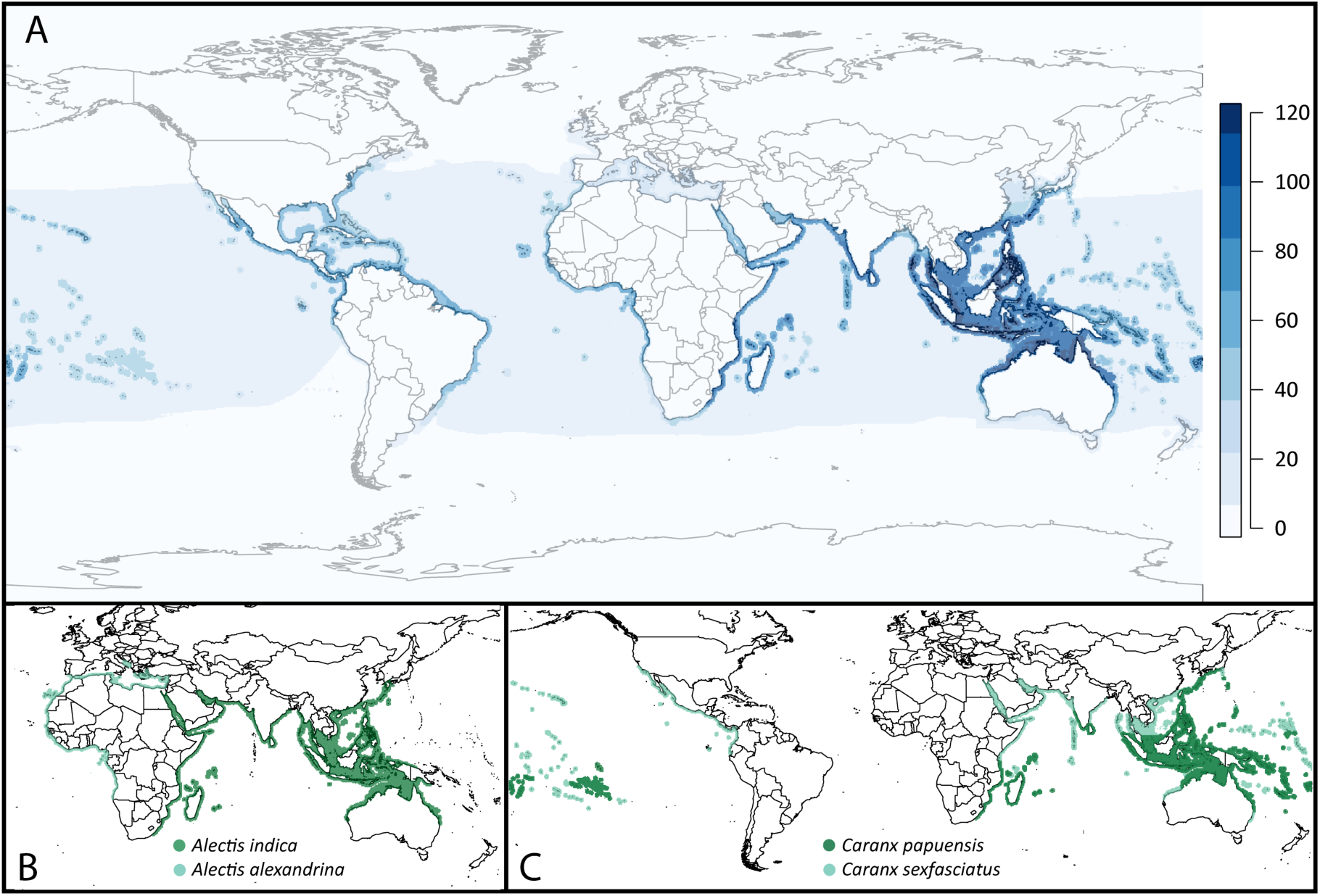
A) Heatmap of Carangoidea species richness using species range extent of occurrence data from IUCN (IUCN 2018) and probability of occurrence data from Aquamaps (Kesner-Reyes et al. 2018). B) Allopatric sister species pair, *Alectis indica* (dark green) and *Alectis alexandrina* (light green), exhibit no range overlap and low range symmetry. C) Sympatric sister species pair, *Caranx sexfasciatus* (light green) and *Caranx papuensis* (dark green), exhibit high range overlap and high range symmetry.

A comprehensive, time-calibrated phylogeny is necessary to study patterns of speciation in Carangoidei. However, the monophyly and taxonomic composition of Carangoidei and the more inclusive lineage, Carangiformes, have been debated since detailed morphological phylogenies were published in the late-20^th^ century (Smith-Vaniz 1984; Bannikov 1986; Gushiken 1988, Girard et al. 2020). Prior to molecular phylogenetic studies, morphological based classifications suggested Carangoidei encompasses the Carangidae, Echeneidae, *Rachycentron canadum*, *Coryphaena*, *Nematistius pectoralis* (Roosterfish), and *Mene maculata* (Moonfish; Smith-Vaniz 1984a,b; Nelson et al. 2016). Molecular data (Harrington et al. 2016; Hughes et al. 2018) and a recent analysis integrating morphological data (Girard et al. 2020), additionally group the morphologically divergent lineages and species *Xiphias gladius* (Swordfish), Istiophoridae (marlins), *Toxotes* (archerfishes) and *Leptobrama* (beach salmon) within Carangiformes. Some molecular studies investigating the relationships among lineages of teleosts that include the major carangiform lineages resolved Carangiformes as paraphyletic (Betancur-R et al. 2013a, 2013b) but most resolve Carangiformes as monophyletic (Near et al. 2013; Harrington et al. 2016; Hughes et al. 2018; Girard et al. 2020). Moreover, molecular phylogenetic analyses of Carangoidei consistently resolve Carangidae as paraphyletic, inclusive of the echenoids, a clade containing *Rachycentron canadum*, *Coryphaena*, and Echeneidae (Reed et al. 2002; Santini and Carnevale 2015; Damerau et al. 2017; Girard et al. 2020).

Although previous phylogenetic studies of Carangoidei disagree on relationships within the clade, these earlier studies are characterized by a combination of limited taxonomic or locus sampling (Reed et al. 2002; Harrington et al. 2016; Damerau et al. 2017) and varying adequacy of fossil calibrations (Santini and Carnevale 2015). Here, we perform a comprehensive phylogenomic analysis using a dataset of more than 955 ultraconserved element (UCE) loci (Alfaro et al. 2018) collected from 80% of the recognized species of Carangoidei. We combine this phylogenetic framework with data on species geographic distributions, depth distributions, and body size to address patterns of allopatry and sympatry in Carangoidei and to examine traits thought to influence speciation by assessing clade-wide patterns of phylogenetic signal and correlation between traits and sister-species contrasts.

## Material and Methods

### Specimen sampling, genomic library construction, and DNA sequencing

We obtained tissues for 154 species through field collection and from multiple museum collections, including nine outgroup species of Carangiformes (Table S1). We extracted DNA using Qiagen DNeasy Tissue kits following manufacturer’s protocol (Qiagen, Inc., Valencia, CA). To prepare DNA samples for library construction, we sheared genomic DNA isolations to obtain a targeted size of 300-600 bp using a sonicator (QSonica Q800R3), and we verified the size distribution using electrophoresis through 1.5% agarose gel. We prepared dual-indexed libraries (Glenn et al. 2019) for targeted enrichment using the KAPA Hyper Prep Library Kit (KAPA Biosystems, Wilmington, MA) following manufacturer’s protocols, although we substituted Sera-mag Speed Beads (Rohland and Reich 2012) for AMPure reagents (Beckman-Coulter, Brea, CA).

We used a probe set designed to target 1314 UCE loci in acanthomorph fishes that is informative for phylogenetic analyses of Carangiformes and other acanthomorph fishes at a variety of evolutionary time scales (Alfaro et al. 2018). We followed library enrichment protocols for the MYcroarray MYBaits kit v3.02 (Arbor Biosciences, Ann Arbor, MI) with three modifications: 1) 100 ng of MYBaits were added to each reaction (a 1:5 dilution of the standard MYBaits concentration), 2) we used 500ng custom blocking oligos designed for our custom index sequences that included 8 inosines to block the 8 nucleotide index sequence, and 3) in place of the human Cot-1 provided by MYcroarray, we used chicken Cot-1 DNA (Applied Genetics Laboratories, Melbourne, FL). Enriched libraries were sequenced using 150 base pair paired-end sequencing on an Illumina HiSeq 4000 (Oklahoma Medical Research Foundation Clinical Genomics Center).

UCE sequence data were processed prior to phylogenetic analyses using the software package *phyluce* v1.6 (Faircloth 2016), which utilizes additional programs to implement each step of the data pipeline. Adapter sequences were trimmed from raw read data using the parallel wrapper (illumiprocessor) to implement the package *trimmomatic* (Bolger et al. 2014). We then assembled cleaned read data into contigs using *Trinity* vr2013-02-25 (Grabherr et al. 2013). To confirm taxonomic identification, we used the *phyluce* program “match_contigs_to_barcodes.py” to extract COI barcode sequence data from assemblies and run a BLAST search on GenBank to verify matches with our DNA sample identification. We constructed alignments of individual UCE loci and performed edge trimming using MAFFT v7.130 (Katoh and Standley 2013). Finally, we generated two data matrices for use in phylogenetic analyses – one for which 75% of taxa were present in each alignment (115 out of 154 taxa) and one for which 95% of taxa were present in each alignment (146 out of 154 taxa).

### Phylogenetic and relaxed molecular clock analyses

We implemented the UCE-specific Sliding Window Site Characteristics approach with site entropy (SWSC-EN) to identify UCE core and flanking regions at each locus (Tagliachollo and Lanfear 2018). We used these results as input for PartitionFinder v2 (Lanfear et al. 2016) to determine the optimal number of partitions for the loci in the 75% complete and 95% complete matrices, using the ‘rclusterf’ search scheme and Bayesian information criteria for model selection.

We inferred a partitioned maximum likelihood (ML) phylogeny using IQ-TREE (Nguyen et al. 2015) and implemented the ultrafast bootstrap approximation approach using 1000 bootstrap replicates and a relaxed hierarchical clustering algorithm (rcluster) that included the top 10% partition merging schemes (Minh et al. 2013). We rooted the tree with the myctophid *Ceratoscopelus warmingii*.

To account for stochasticity in the evolutionary history amongst individual UCE loci, we performed a coalescent-based analysis using loci from the 75% complete matrix. We first generated individual locus trees in MrBayes v3.2.6 using MCMC chains run for 1 million generations (model HKY + gamma), sampling trees every 200 generations and discarding the first 25% of samples as burnin before summarizing the posterior sample of trees as a 50% majority rule consensus tree (Ronquist et al. 2012). We inferred a species tree from these locus trees using ASTRAL-II v5.6.2 using the default parameters (Mirarab and Warnow 2015).

We used a relaxed molecular clock approach in BEAST v1.10.4 (Suchard et al. 2018) to estimate divergence times of carangiform lineages. Because BEAST has computational limitations when analyzing hundreds of loci simultaneously, we performed replicate analyses using different combinations of loci, *sensu* Harrington et al. (2016) and Brantstetter et al. (2017). We generated 4 random subsets of 25 loci from the 371 loci in the 95% complete matrix. For each subset of 25 loci, we used the SWSC-EN approach to UCE partitioning as input for PartitionFinder to identify optimal partitioning schemes for each BEAST analysis using Bayesian Information Criteria (Lanfear et al. 2016; Tagliacollo and Lanfear 2018). We applied an uncorrelated lognormal clock model in BEAST with a GTRGAMMA site substitution model and a birth-death speciation tree model, and we fixed the topology to the IQ-TREE-inferred tree from the 75% complete matrix. We included nine fossil calibration points from Harrington et al. (2016) that spanned the Carangiformes clade and assigned the same lognormal prior distributions to incorporate age priors for select nodes. To assess convergence of the model parameters, we ran 15 analyses of each of the 4 subsets of randomly chosen 25 loci for 100 million generations and used Tracer v1.70 (Rambaut et al. 2018) to determine convergence and sufficient mixing of the MCMC chain (we ensured ESS > 200). We assigned a burn-in value of 25 million generations. For each of the 4 subsets, we used LogCombiner v1.10 to combine individual analyses and TreeAnnotator v1.10 to construct a maximum clade credibility tree (Drummond et al. 2012). Summarized node heights were rescaled in TreeAnnotator to reflect posterior median heights.

### Biogeographic and trait analyses

We obtained range data for 125 species from the IUCN Red List database, which consists of range maps, validated by experts, depicting the known “extent of occurrence” of each species in the form of spatial polygons (IUCN 2018). We obtained ranges for species missing from the IUCN database (N = 25) from Aquamaps, a database of species range predictions that uses a combination of occurrence data and species range modeling to assign relative probabilities of occurrence for each point coordinate (Kaschner et al. 2016). To make the two datasets comparable, we interpreted any non-zero probability of occurrence from the Aquamaps database as an occurrence point, which essentially makes that species’ range an extent of occurrence, equivalent to the IUCN database (O’Hara et al. 2017). Species for which no range data were available (N = 4) were trimmed from the BEAST-inferred time tree prior to biogeographic analyses. We also trimmed the BEAST-inferred time tree to only include carangoid taxa (Carangidae, Echeneidae, Coryphaenidae and Rachycentridae). The final dataset contained 123 species (Table S1). We rasterized IUCN shapefiles and combined them with the Aquamaps dataset at the same grid and resolution (0.5° x 0.5°) using the Behrmann equal-area cylindrical projection. Geographical range for a given species was defined as the sum of all polygons in that species’ distribution map. All spatial analyses were performed using GDAL and R v3.5.2 (R Core Team 2019).

To assess the relationships between morphological traits, ecological traits, and biogeography, we compiled trait data on maximum body length and maximum depth in the water column for the 125 carangoid species from the IUCN database (IUCN 2018). If IUCN data were missing, we used FishBase (Froese and Pauly 2016). We also compiled data on habitat class (reef or non-reef-associated) and diet (piscivorous or non-piscivorous) from a prior study (Frédérich et al. 2016) and the IUCN database (IUCN 2018).

We extracted divergence time estimates for each node of the time-calibrated phylogeny and conducted pairwise comparisons of range overlap and range symmetry *sensu* Chesser and Zink (1994) using custom scripts in R v3.5.2, as well as the packages ‘ape’ (Paradis and Schliep 2019), ‘geiger’ (Pennell et al. 2014), ‘phytools’ (Revell 2012), and ‘picante’ (Kembel et al. 2010). We defined range overlap as the area occupied by a given species pair divided by the area of the species with a smaller range (Chesser and Zink 1994). This produces an index ranging from 0 to 1, with 0 indicating no overlap (Fig. 1B) and 1 indicating complete overlap (Fig. 1C). Complete overlap meant that both species co-occur throughout their entire respective ranges or that the range of one species is entirely encompassed by the other. We classified species as allopatric if the range overlap index was less than 0.05 and sympatric if the range overlap index was any value greater than or equal to 0.05. We performed these analyses between all species in the phylogeny and extracted sister species pairs for additional analyses. A sister species pair was defined as two species sharing a unique common ancestor, i.e., an ancestor not shared with any other taxa in our phylogeny. We analyzed 41 distinct sister species pairs, of which at least 32 we believe to be direct sister species (reciprocally monophyletic) based on molecular data. The remaining nine pairs out of 41 were uncertain due to unsampled species which may represent a closer relative to one of the species in those pairs. Throughout the rest of the manuscript, we refer to sister species pairs inclusive of those believed to be direct sister species and those that exclusively share a single ancestor based on our phylogenetic sampling. To test for peripatry, we calculated range symmetry for each sister species pair, which we defined as the smaller of the two species’ ranges divided by the sum of both species’ ranges (Barraclough and Vogler 2000). The range symmetry metric falls between 0 and 0.5, where 0.5 indicates that both species have equal-sized ranges.

### Phylogenetic signal in Carangoidei

To examine the extent to which closely related species in Carangoidei were ecologically and physically similar, we tested for phylogenetic signal, defined as the tendency for related species to resemble each other more than they resemble species drawn at random from the tree (Hillis and Huelsenbeck 1992; Blomberg et al. 2003). We tested three continuous traits (body length, water column depth and range size) and two discrete traits (habitat and piscivory). For the continuous traits, we tested for phylogenetic signal using Blomberg’s *K* (Blomberg et al. 2003) implemented in the R package ‘phytools’ (Revell 2012). A value of Blomberg’s *K* equal to one indicates that the variance in trait values exhibited by species are distributed across the phylogeny in a manner consistent with Brownian motion; *K* values lower than one imply less variance amongst species’ trait values than expected by a Brownian process, while *K* values greater than one indicate that the variance of species’ traits is more distributed among clades, rather than within clades (i.e., a strong relationship between traits and relatedness; Blomberg et al. 2003). After visually assessing the distribution of trait data, we normalized the variables for body length and water column depth by performing a log-transformation. Blomberg’s *K* was significant (p < 0.05) for all continuous variables: body length, water column depth, and range size. However, since *K* only compares expectations to a Brownian motion model, which has no constraints on evolution, we tested support for other models of evolution (Ornstein-Uhlenbeck and Early- burst) to determine the most appropriate model when performing phylogenetic least squares regression. We determined the best-supported model of evolution by ranking sample-size corrected Akaike information criterion (AIC) values (Hurvich and Tsai 1986). The Ornstein- Uhlenbeck (OU) model was favored over the Brownian motion and Early-burst models (Table S2); therefore, we examined correlations between body length and water depth and body length and range size using phylogenetic least squares regression with an OU error model using the package ‘nlme’ in R (Pinheiro et al. 2020). We used Martins’ and Hansen’s correlation structure (Martins and Hansen 1997) and an intraspecific standard error value (0.0772) generated from 49 clades of vertebrate organisms (Harmon et al. 2010).

We tested for phylogenetic signal in the discrete traits – habitat (reef or non-reef) and piscivory (piscivorous or non-piscivorous) – using Fritz’s *D* (Fritz and Purvis 2010) in the R package ‘caper’ (Orme et al. 2012). The value of Fritz’s *D*, which ranges from negative one to one, is equal to one if the distribution of binary traits, accounting for phylogeny, is random. Fritz’s *D* is zero if the distribution of traits resembles what is expected under a Brownian motion model, while Fritz’s *D* is below zero if the distribution of traits is more phylogenetically conserved than expected under Brownian motion (e.g., traits are clustered on a phylogeny).

We tested for phylogenetic signal of range overlap and range symmetry in Carangoidei using multiple matrix regression by means of a partial Mantel test with 1000 phylogenetically informed permutations using ‘phytools’ in R (Mantel 1967; Revell 2012). Although the Mantel test has been criticized for its low power, it is a suitable option for testing phylogenetic signal in data that are inherently pairwise contrasts, such as measures of range overlap and symmetry (Harmon and Glor 2010).

We used linear regression to examine the relationships between divergence times for 41 sister species pairs, range overlap, and range symmetry because these metrics are hypothesized to be informative about speciation mechanisms (Chesser and Zink 1994; Barraclough and Vogler 2000; Anacker and Strauss 2014). We also used Welch’s *t*-tests to statistically examine the effect of allopatric and sympatric sister species pairs on ecological traits. For each sister species pair, we calculated trait contrasts, specifically the differences in body length and maximum water column depth between sister species. We also analyzed differences in body length and water column depth for sympatric sister species categorized by habitat type, i.e., whether both sister species occupied the same or different habitat type (reef or non-reef). We excluded one sister pair, *Trachinotus mookalee* and *Trachinotus anak*, from depth analyses because no maximum depth data are currently available for *T. mookalee* (Froese and Pauly 2016; IUCN 2018).

## Results

### Phylogenomic analyses and divergence times

We collected sequence data from an average of 958 loci for 154 individuals. Following alignment trimming, mean locus length was 972 bp (range: 319 – 1570 bp) and each locus contained a mean of 270.6 parsimony informative sites. The 75% complete alignment included 986 loci and the 95% complete alignment included 371 loci. PartitionFinder produced 143 partitions for the 75% complete matrix and 94 partitions for the 95% complete matrix. IQ-TREE (Fig. S1, S2) and ASTRAL (Fig. S3) inferred nearly identical trees with high bootstrap support and local posterior probabilities, respectively (Fig. S4, S5). Given the near-identical topologies generated from the 75% complete concatenated dataset compared to the coalescent-based tree (Fig. S4), we used the 75% complete concatenated IQ-TREE as a fixed topology in BEAST for divergence time estimation (Fig. 2).

**Figure 2.**
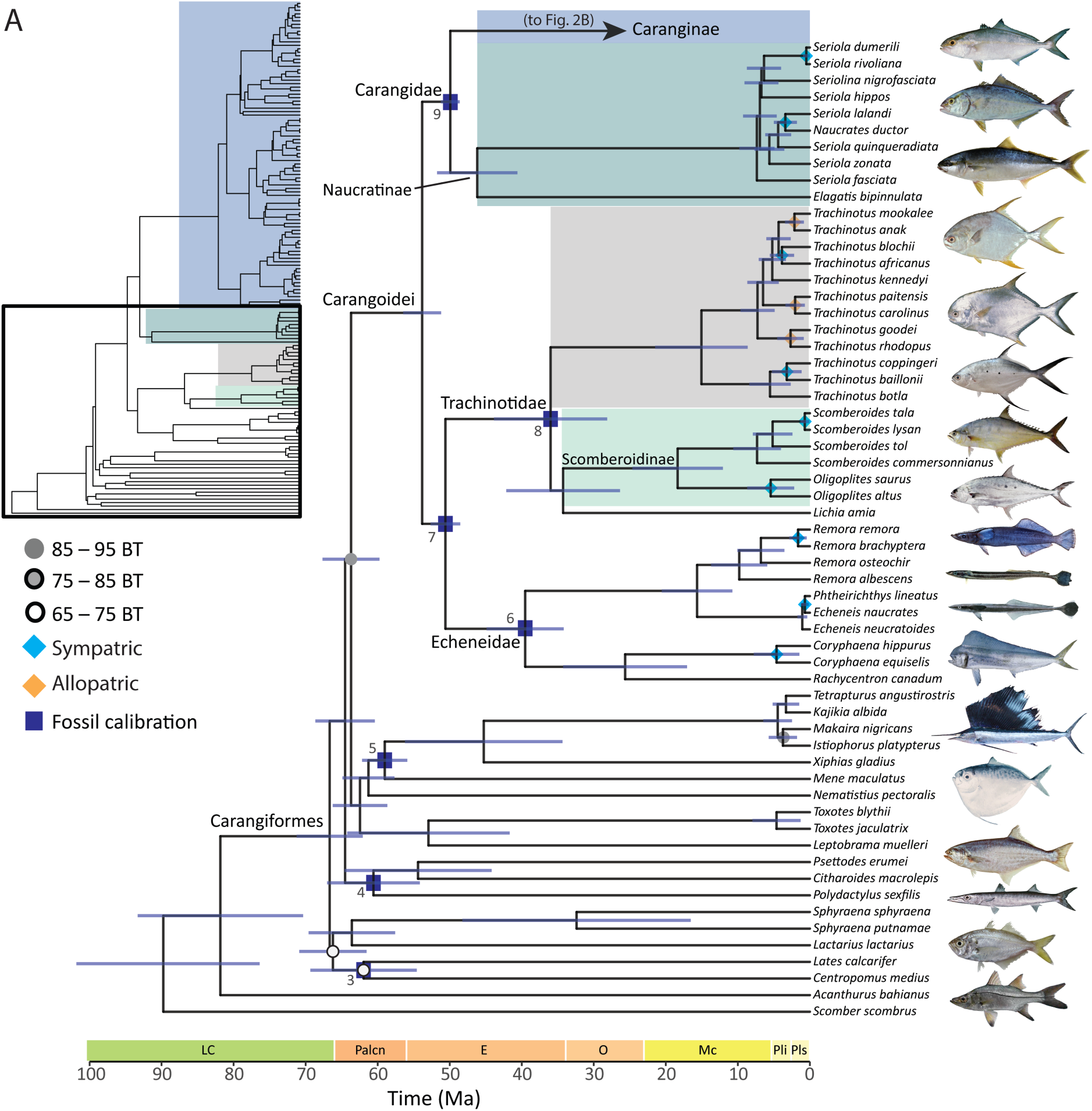
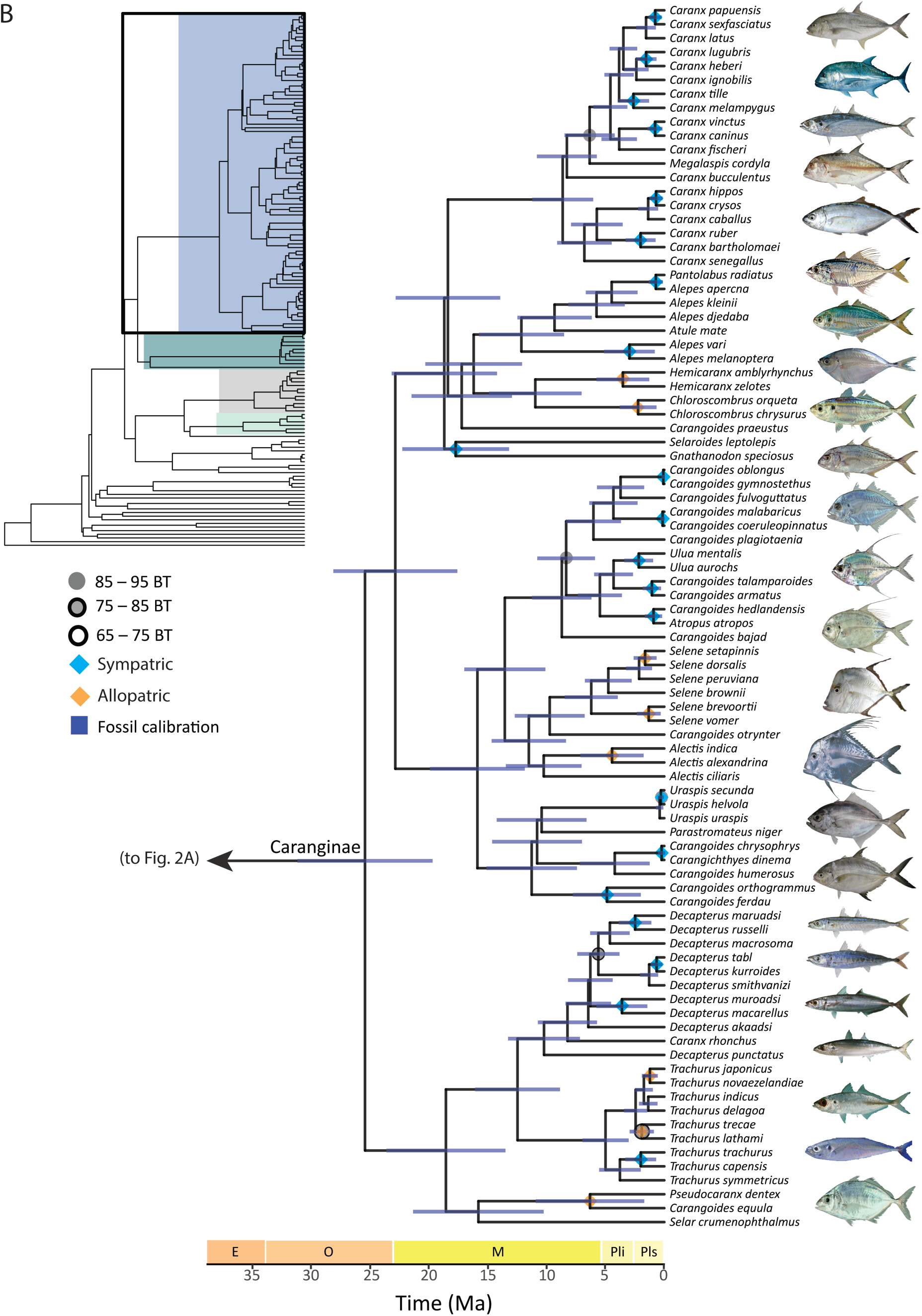
Time-calibrated phylogeny of 145 species of Carangiformes and two outgroup species generated by BEAST using a guide tree from the 75% complete matrix constructed in IQ-TREE and nine fossil calibration points. Nodes represent median ages from a maximum clade-credibility tree. Bootstrap support values are indicated as circles on each node. No circle indicates 95-100% BT support (BT). Dark blue rectangles indicate nodes calibrated with priors based on fossil data. Diamond node labels indicate sympatric (light blue) and allopatric (black) sister species pairs. Fish image sources are located in Table S4.

The majority of nodes in the IQ-TREE-inferred phylogeny generated from the 75% complete matrix are strongly supported; 90% of nodes have bootstrap support values of 100 (Fig. 2). Our phylogenomic analyses of the UCE loci suggest four distinct lineages within carangiform fishes: 1) a clade containing *Lates calcarifer*, Centropomidae, *Lactarius lactarius* and *Sphyraena*, 2) a clade containing Polynemidae and Pleuronectoidei (flatfishes), 3) a clade containing *Leptobrama*, *Toxotes*, *Nematistius pectoralis*, *Mene maculata*, *Xiphias gladius*, and Istiophoridae, and 4) a clade containing Echeneidae, *Rachycentron canadum*, and *Coryphaena* nested in a paraphyletic Carangidae (Fig. 2A, S1 – S3).

Within Carangoidei, our phylogenetic hypotheses inferred from UCE data resolve two large subclades. The first consists of the echenoids (Echeneidae, *Rachycentron canadum*, and *Coryphaena*) which we reclassify as Echeneidae *sensu* Rafinesque (1810). This clade is inferred as the sister lineage of a clade we elevate to Trachinotidae containing *Trachinotus*, *Lichia amia* and Scomberoidinae (*Oligoplites* and *Scomberoides*). *Lichia amia* was previously classified with *Trachinotus* in Trachinotini (Smith-Vaniz 1984; Nelson et al. 2016); however, the UCE data infer *L. amia* as the sister lineage of Scomberoidinae with strong node support (Fig. 2, S1 – S3). We delimit the second major subclade within Carangoidei as Carangidae, inclusive of Naucratinae (*Elegatis*, *Naucrates*, *Seriola*, and *Seriolina*) and Caranginae (*Alectis, Alepes, Atropus, Atule, Carangoides, Carangichthys, Caranx, Chloroscombrus, Decapterus, Gnathanodon, Hemicaranx, Megalaspis, Pantolabus, Parastromateus, Pseudocaranx, Selar, Selaroides, Selene, Trachurus, Ulua, Urapsis*; Fig. 2). Naucratinae and Caranginae contain numerous paraphyletic genera. For *Alepes*, *Decapterus*, *Seriola*, and *Caranx,* paraphyly is the result of one or two species classified in other genera resolving within the clade (Fig. 2, S1 – S3). *Carangoides* is polyphyletic, with species distributed across nine clades (Fig. 2B, S1 – S3). Our phylogenetic hypotheses inferred from UCE data are broadly congruent with recent molecular analyses focusing on Carangoidei that contain dense taxonomic sampling (Santini and Carnevale 2015; Harrington et al. 2016; Damerau et al. 2017).

Within Carangiformes, species relationships inferred from the UCE data are largely consistent across the different methods of analysis (IQ-TREE, ASTRAL) and matrix composition (75% or 95% complete). Differences in phylogenetic relationships between the 75% IQ-TREE-inferred topology and ASTRAL coalescent-based tree involve the phylogenetic placement of *Centropomus medius*, *Seriola nigrofasciata*, *Decapterus macarellus and D. akaadsi, Caranx crysos* and *C. caballus*, and *Parastromateus niger* (Fig. S4). Phylogenies inferred using IQ-TREE from the 75% and 95% UCE matrices differ in the resolution of *Trachurus trecae*, *Decapterus akaadsi*, *Uraspis uraspis* and *Carangoides bajad* (Fig. S5).

Using relaxed-clock molecular dating analyses, we generated similar estimates of divergence times across the four random subsets of 25 UCE loci. We present specific date estimates and branch lengths from one subset of 25 UCE loci (Fig. 2), but the four subsets produced similar node ages and overlapping 95% highest posterior densities (HPD; Fig. S6). These analyses estimate the age of the most recent common ancestor (MRCA) of Carangiformes as 66.66 Ma (95% HPD: 62.43 – 71.59 Ma) and of Carangoidei as 53.82 Ma (95% HPD: 51.50 – 56.77 Ma; Fig. 2). These divergence times are younger than a previous study on Carangoidei (Santini and Carnevale 2015) but fall within the 95% confidence intervals of other phylogenetic studies that estimate the ages of Carangiformes and Carangoidei (Near et al. 2013; Harrington et al. 2016; Alfaro et al. 2018; Hughes et al. 2018).

### Phylogenetic signal

Tests of continuous trait variables within Carangoidei using Blomberg’s *K* suggest there is phylogenetic signal for body length (*K* = 0.123, *p* = 0.001), water column depth (*K* = 0.090, *p* = 0.006) and range size (*K* = 0.307, *p* = 0.001). Phylogenetic least squares regression with an OU error model suggests a significant correlation between maximum body length and maximum water column depth (*t* = 2.593, *p* = 0.011), but not between maximum body length and geographic range size (*t* = 1.004, *p* = 0.317).

We calculated Fritz’s *D* to examine phylogenetic signal in discrete variables (reef habitat and piscivory) and compared observed *D* values to simulated sums of expected character changes under Brownian motion and random models. The test for reef habitat (*D* = 0.663) suggests a significant departure from Brownian motion expectations (*p*[*D* > 0] < 0.001) but more phylogenetic signal than expected from a random distribution of habitat traits across the phylogeny (*p*[*D* < 1] < 0.001). The test of Fritz’s *D* for diet (piscivory or non-piscivory; *D* = 0.069) suggests the evolution of diet resembles a Brownian process (*p*[*D* > 0] = 0.365) rather than a random process (*p*[*D* < 1] < 0.001). Mantel tests of contrast variables reveal a significant correlation between phylogeny and overlap of geographic ranges (R^2^ = 0.020, *p* = 0.009), as well as phylogeny and geographic range size symmetry (R^2^ = 0.033, *p* = 0.001).

### Sister species analyses

Among the 41 resolved carangoid sister species pairs, 30 (73%) are sympatric (range overlap > 0.05) and 11 (27%) are allopatric (Fig. 2). All allopatric sister pairs have range overlap values of zero except one pair, *Trachinotus anak* and *T. mookalee*, which has a range overlap value of 0.02. All sympatric species pairs have range overlap values > 0.6 except one, *Uraspis helvola* and *U. secunda*, whose range overlap is 0.16 (Fig. 3A). The node ages of sympatric sister species pairs range from 0.08 – 17.74 Ma, with a median age of 1.65 Ma, while the node ages of allopatric pairs range from 1.23 – 6.31 Ma, with a median age of 2.14 Ma. We find no effect of node age on range overlap (*r* = 2.25 x 10^-4^, *p* = 0.926; Fig 3A) or geographic range size symmetry (*r* = 0.013, *p* = 0.480), nor is there a significant correlation between range overlap and range size symmetry (*r* = 0.072, *p* = 0.091; Fig. 3B). Median range size symmetry is 0.273 for allopatric species pairs and 0.348 for sympatric species pairs. Notably, there are greater differences in maximum water depth between sister species in sympatry versus those in allopatry (*t* = 2.513, *p* = 0.017; Fig. 4A). We also observe greater differences in maximum body length between sympatric sister species pairs compared to allopatric pairs (Fig. 4B), though these are not significant (*t* = 1.821, *p* = 0.081). After excluding four sister species pairs that were not sufficiently resolved by the ASTRAL coalescent-based tree topology, we examined differences in water column depth and body length between the remaining 37 pairs, of which 10 were allopatric and 27 sympatric. With this reduced dataset, we again observe high sympatry (73%) and greater differences in water column depth (*t* = 2.028, *p* = 0.054) and body length (*t* = 2.377, *p* = 0.026) in sympatric pairs compared to allopatric pairs (Fig. S7).

**Figure 3.**
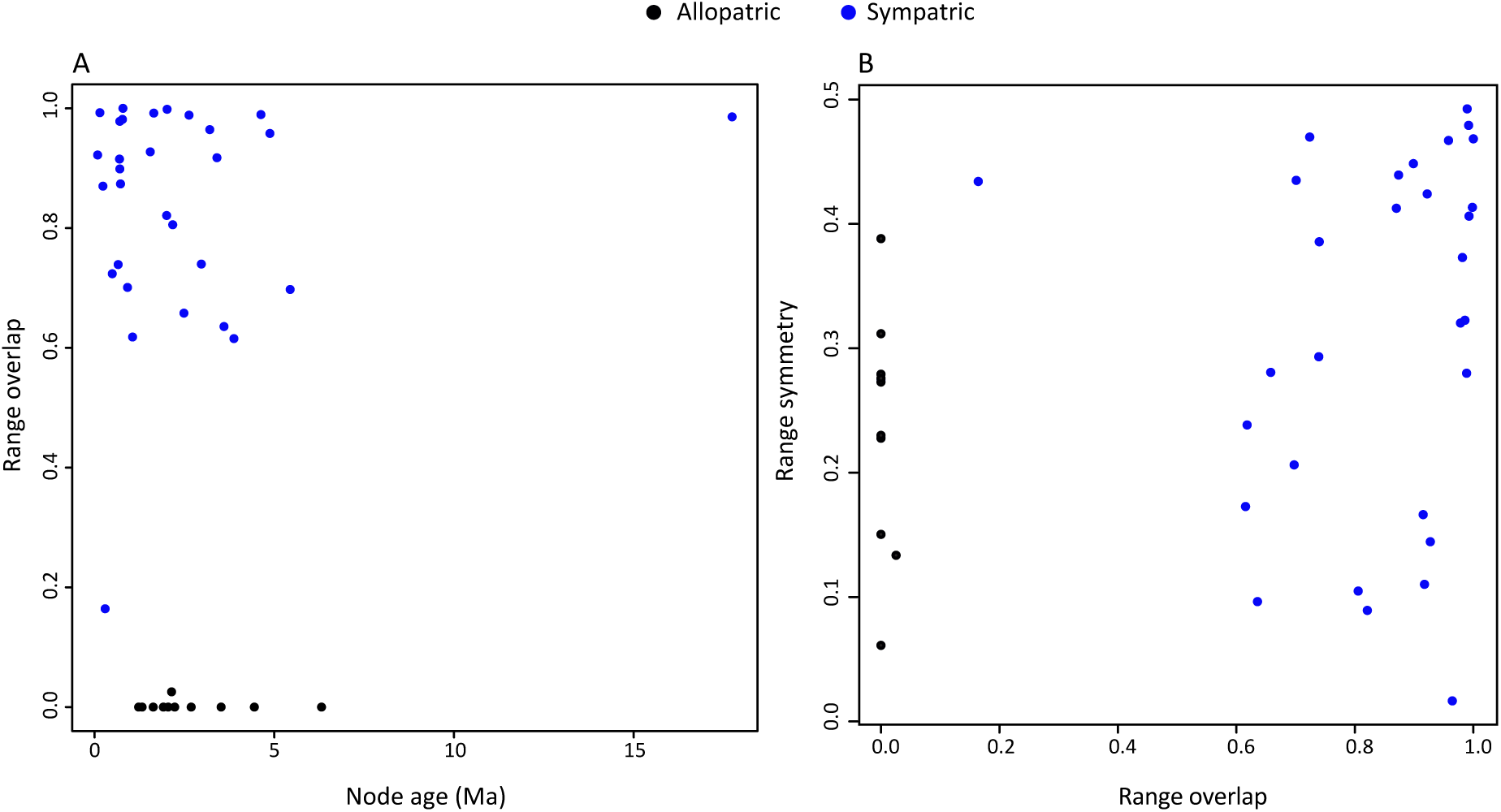
Range overlap as a function of node age (A) and range symmetry as a function of overlap (B) for 41 sister species pairs within Carangoidea.

**Figure 4.**
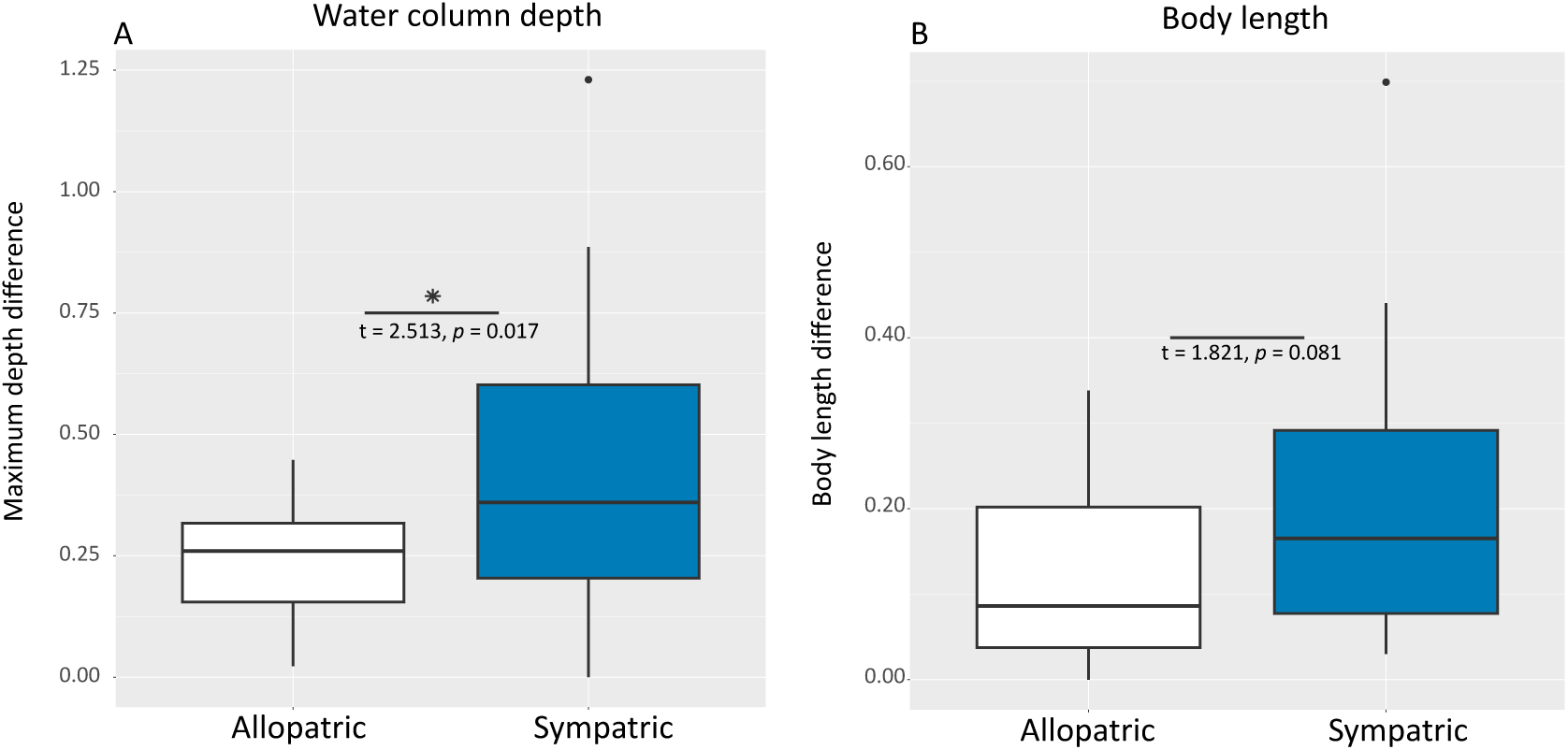
Contrasts between allopatric and sympatric sister species pairs for maximum water column depth (A) and maximum body length (B). Results of Welch’s *t-*tests are presented to show significance between allopatric and sympatric sister species pairs.

Out of 11 allopatric pairs from the 41-pair dataset, 73% (N = 8) contain two non-reef-associated species, whereas 40% of sympatric pairs (N = 12) contain two reef-associated species, 40% (N = 12) contain one reef and one non-reef-associated species, and 20% (N = 6) contain two non-reef-associated species (Table S3). We find greater differences in maximum water depth (*t* = 2.173, *p* = 0.034) between sympatric sister species pairs that occupy the same habitat (e.g., both occupy reef or non-reef habitats) compared to sympatric pairs that occupy different habitats (Fig. 5).

**Figure 5.**
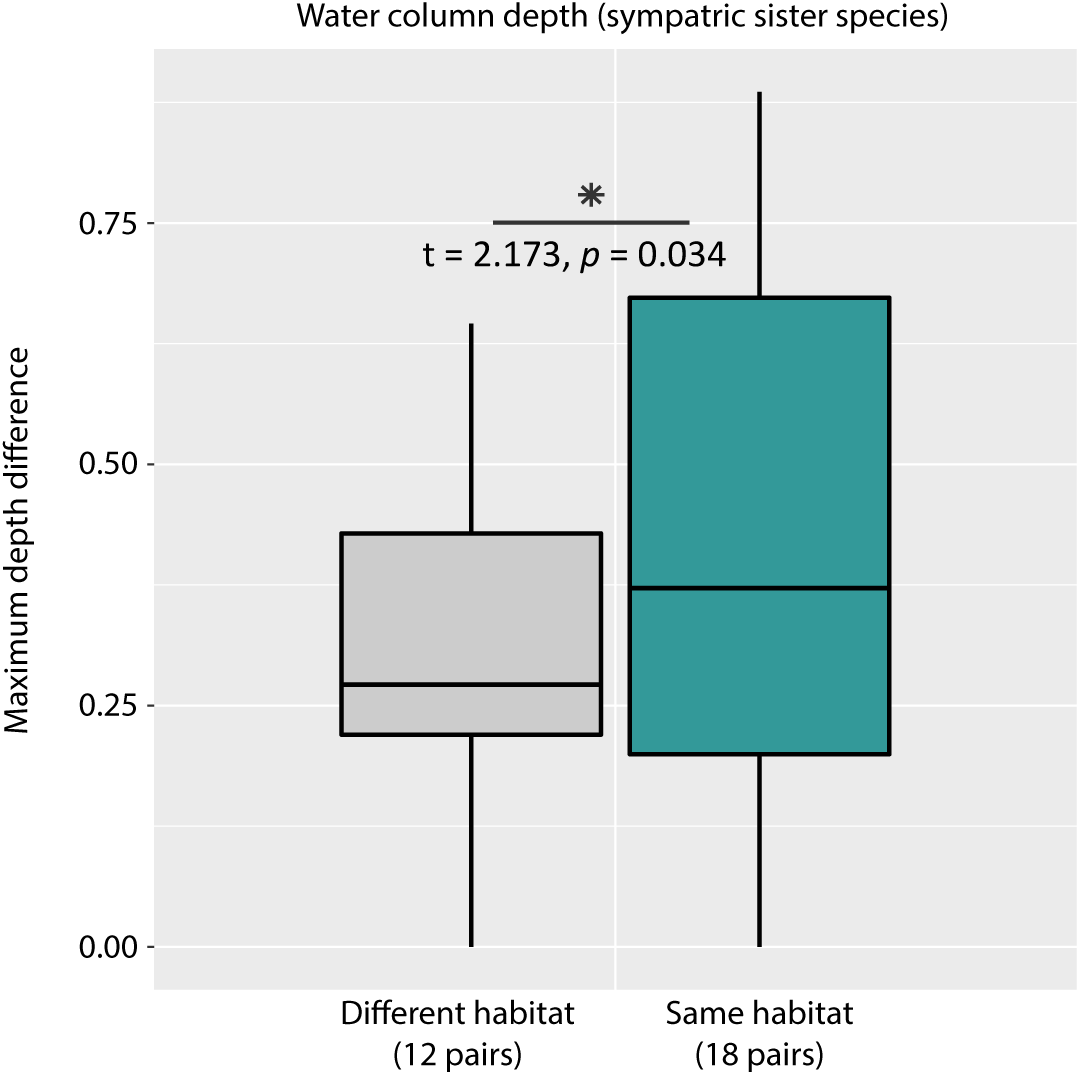
Contrasts between sympatric sister species pairs for maximum water column depth based on similarity in habitat type, i.e., whether both species occupy the same habitat (both reef or both non-reef) or different habitats. Results of Welch’s *t-*tests are presented to show significance between habitat types.

## Discussion

### Carangiform phylogeny and timing of diversification

With a dataset averaging 958 UCE loci and representing 80% of the known species diversity within Carangoidei, we provide strong phylogenomic resolution for the relationships within this clade. The molecular and phylogenomic perspective on carangoid relationships is notable in the consistent paraphyly of the traditional delimitation of Carangidae when excluding the echenoids; thus, we present a new classification for the carangoid subclades Carangidae, Echeneidae and Trachinotidae. Our resolution of subclades within Carangiformes is concordant with previous analyses using UCEs, with the exception of the relationships of Latidae, Centropomidae, and *Sphyraena* (Harrington et al. 2016; Girard et al. 2020); these different phylogenetic relationships of early diverging carangiform lineages reflect ongoing challenges using molecular data to resolve taxonomic relationships due to short internal branches (Felsenstein 1978; Kubatko and Degnan 2007).

Our estimates of node ages suggest the origin of Carangoidei was ∼53 Ma in the early Eocene (Fig. S6). Estimates from UCE data for the age of Carangoidei are much younger than a previous study which suggested a Late Cretaceous origin; this is likely due to that study’s fossil calibrations, which have older age estimates within Carangoidei (Santini and Carnevale 2015). Due to several identical fossil calibration points within Carangiformes shared across studies, our age estimates are similar to phylogenomic analyses using UCEs, (Harrington et al. 2016; Alfaro et al. 2018) and exons from protein coding genes (Hughes et al. 2018). Our results suggest most of the species level diversification occurred during the last 10 million years, in the late Miocene (∼11.63 – 5.33 Ma). The late Miocene was a period of warmer global climate and expanding coral reef habitats, which is congruent with the observed diversification of other tropical and sub-tropical coral reef fish lineages with diversity centered in the Indo-Pacific Archipelago region (Cowman and Bellwood 2013; Cowman 2014). This diversification may have been driven by the formation of coral reef habitats in the Indo-Pacific Archipelago after the closing of the Tethys seaway towards the end of the Miocene, ∼5.33 Ma (Rogl 1999; Cowman and Bellwood 2013).

### Patterns of carangoid sympatry and allopatry

While sympatry of sister species pairs is ubiquitous across Carangoidei, regardless of node age, 27% of sister species pairs were allopatric. Most cases of allopatry (64%) were likely caused by vicariance, specifically the rise of the Isthmus of Panama. Seven out of eleven allopatric pairs have divergence times younger than 5 Ma and presently exhibit parallel range patterns, where one species occupies the eastern Pacific (e.g., *Selene brevoortii*) and its sister species inhabits the western Atlantic (e.g., *Selene vomer;* Fig. 6A). The other cases of allopatry are potentially maintained by the cold-water barrier caused by the Benguela and Agulhas currents off the southern coast of South Africa separating the Atlantic and Indian Oceans (*Alectis indica* and *A. alexandrina*), or open-ocean barriers in the Atlantic (e.g., *Selene setapinnis* and *S. dorsalis*; Fig. 6B) and Indo-Pacific (*Trachinotus mookalee* and *T. anak*; *Trachurus japonicus* and *T. novaezelandiae*). The isolating barriers are unknown for the oldest (∼6.3 Ma) diverging allopatric sister pair, *Pseudocaranx dentex* and *Carangoides equula*, which are found throughout the Atlantic and Indo-Pacific, respectively.

**Figure 6.**
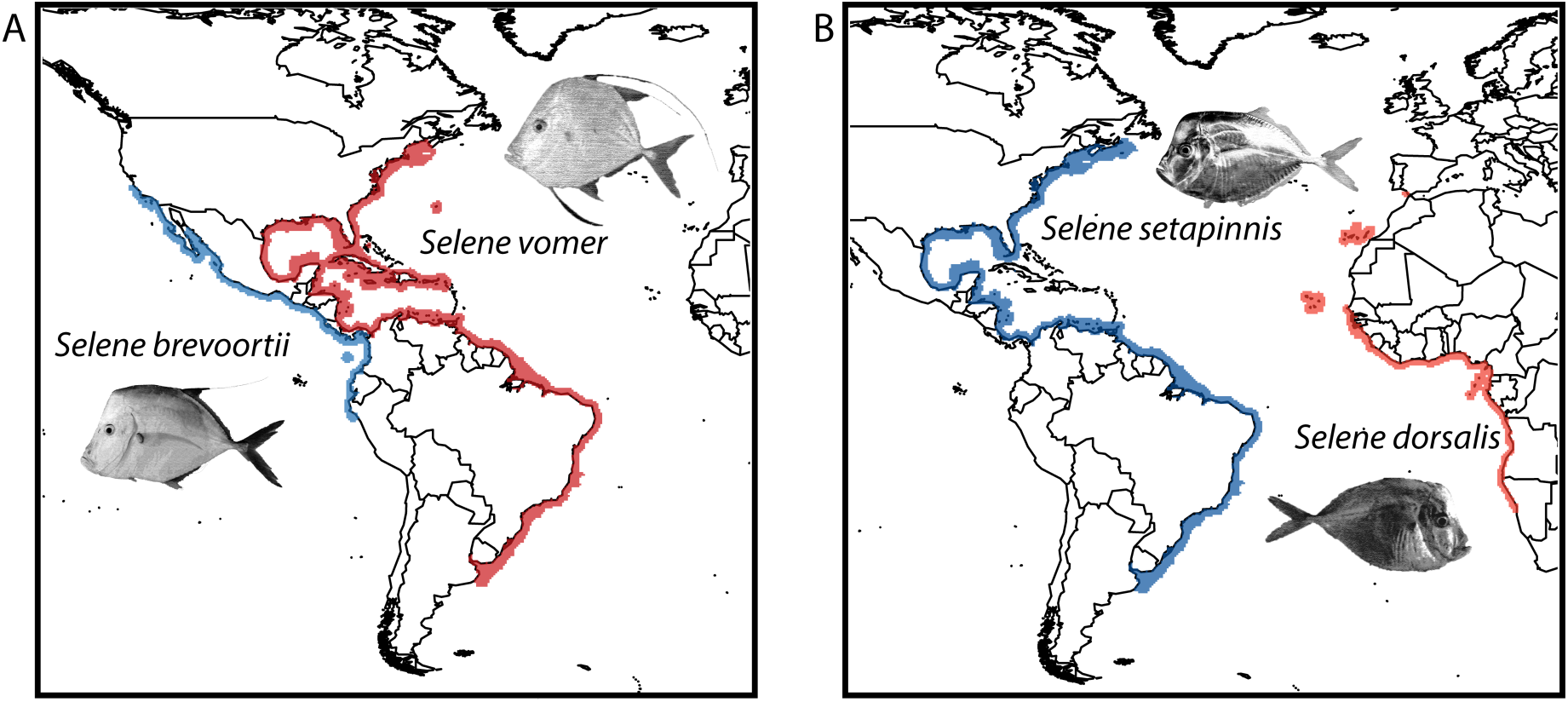
Examples of allopatric sister species pairs showing (A) the most prevalent scenario of two sister species separated by the Isthmus of Panama, demonstrated by *Selene brevoortii* (blue) and *Selene vomer* (red), and (B) an example of an open ocean barrier in the Atlantic Ocean separating sister species *Selene setapinnis* (blue) and *Selene dorsalis* (red).

No study, to our knowledge, has demonstrated such widespread sympatry in a large lineage of marine fishes, although high degrees of sympatry have been demonstrated in some genera of coral reef fishes – for example, over 80% of sister species pairs in *Pomacanthus* angelfishes (N = 13 spp.; Hodge et al. 2013) and *Haemulon* grunts (N = 21 spp.; Rocha et al. 2008). Most comparable analyses of marine fishes have found a higher prevalence of allopatry between sister species, from 62% in New World haemulid fishes (N = 42 spp.; Tavera and Wainwright 2019), to 64% in parrotfishes (N = 61 spp.; Choat et al. 2012), to 88% in *Halichoeres* wrasses (N = 24 spp.; Wainwright et al. 2018). In *Holocanthus* angelfishes (N = 7 spp.), one sister pair is sympatric while the other species are allopatric (Tariel et al. 2016).

Following long-held assumptions that allopatric speciation is the predominant mechanism driving species diversification (Turelli et al. 2001; Coyne and Orr 2004), the clade-wide pattern we observed suggests secondary sympatry occurring after allopatric speciation. Moreover, the positive association between range overlap and range symmetry implies close relatives can co-exist and maintain sympatry across large portions of their ranges. The greater divergence in body size and water column depth in sympatric sister pairs compared to allopatric pairs may be one mechanism that reduces interspecific competition, facilitating secondary sympatry among closely related species. Similar examples of transitions from allopatry to secondary sympatry are sparse but have been observed in birds (Phillimore et al. 2008; Martin et al. 2010; Andersen et al. 2015) and coral reef fishes (Quenouille et al. 2011; Pigot and Tobias 2015). Reef fishes and marine cetaceans exhibit higher transition rates to sympatry than birds and other vertebrate lineages (Pigot and Tobias 2015). Although node age is a significant predictor of transition from allopatry to sympatry in terrestrial organisms, the probability of sympatry is independent of node age in coral reef fishes and cetaceans due to frequent, fast transitions between allopatric and sympatric states (Pigot and Tobias 2015). While these authors attribute the fast transitions to higher intrinsic dispersal abilities in lineages of marine organisms compared to terrestrial vertebrates (Pigot and Tobias 2015), it has been shown that dispersal ability - including pelagic larval duration - has a nuanced correlation with range size (Lester et al. 2007; Mora et al. 2012).

### Ecological signature of secondary sympatry in carangoid fishes

We observe higher divergence in maximum water depth and body size in sympatric sister species pairs, suggesting ecological factors facilitated sympatry among the most closely related species of carangoids, which have maintained genetic isolation over time. Water depth differences between sympatric sister species have also been documented in New World *Halichoeres* fishes (Wainwright et al. 2018), and sympatric sister pairs of parrotfish exhibit greater differences in body size, morphology, habitat type and color patterns (Choat et al. 2012). Body size and water column depth are reflective of resource use, with the former being a strong correlate of prey consumption (Romanuk et al. 2011) and the latter indicative of ecological niche partitioning (Platell and Potter 2001; Ingram 2011). The trait differences we examined in Carangoidei may be the result of character displacement, which is represented by the divergence of character traits in two or more lineages occurring in sympatry (Schluter and McPhail 2002; Brown and Wilson 2006; Pfennig and Pfennig 2010; Pigot and Tobias 2013), but at present, character displacement is difficult to prove due to a lack of detailed ecological trait data across carangoid species’ ranges. Given that body size and maximum depth in the water column are positively correlated (Smith and Brown 2002), it is unclear if divergence in body size is driving divergence in water depth distribution. Further research on these and other traits is warranted, particularly to compare sister species’ traits between areas of overlap versus non-overlap.

The displacement of ecological and behavioral characters, in part by minimizing competition, is hypothesized to facilitate sympatry between closely related species (Brown and Wilson 2006; Pfennig and Pfennig 2010). In coral reef environments, habitat complexity may influence character displacement in reef fishes, be it divergence in mate recognition (Hemingson et al. 2019), trophic partitioning (Price et al. 2013), reef preference (Choat et al. 2012), or territoriality (Nursall 1974; Fricke 1980). Yet, since fewer than half of carangoid species (45%) are classified as reef-associated, carangoid niche partitioning might be shaped by different factors than those affecting coral reef fishes. Most (80%) allopatric carangoid sister pairs are non-reef-associated, while 80% of sympatric sister species contain at least one reef-associated species. A previous analysis on carangoid body shape and ecological traits found that shifts from reef to non-reef environments increased rates of morphological diversification, implying that non-reef environments influenced morphological changes more than reef environments (Frédérich et al. 2016). Although the authors did not find an effect of habitat type on rates of phylogenetic lineage diversification, this could be due to their age estimates of Carangoidei (Santini and Carnevale 2016), which are substantially older than our age estimates and those of other phylogenomic studies (Harrington et al. 2016; Hughes et al. 2018). Our results suggest habitat and diet resemble a Brownian motion model of trait evolution, but we did not test for the effects of trait evolution on rates of diversification.

Ecological partitioning among closely related species occupying non-reef environments might be one reason why Carangoids exhibit such high disparity in body shape and body size relative to other percomorphs (Gushiken 1988; Price et al. 2015). Despite this variation in body shape and size, we still observe phylogenetic signal in body length, which corroborates previous morphological work suggesting similarity in the evolution of Carangoid body types amongst major subclades (Reed et al. 2002). Our tests of phylogenetic covariance suggest that the evolution of certain morphological and ecological traits has been conserved during carangoid lineage diversification. Notably, though we observe phylogenetical signal in body length and water column depth in Carangoidei, these are divergent traits between sister species pairs, with greater contrasts between sympatric species. Our results highlight the benefits of performing sister species analyses, not only because such analyses pose less risk of overestimating divergence times due to extinction events (Hodge and Bellwood 2015), but also because they are independent replicates that are less likely to be phylogenetically confounded (Felsenstein 1985) and may reveal trait divergences that are masked by analyses of phylogenetic signal across the entire clade.

## Conclusions

Analyzing sister species pairs, we found strong evidence for widespread sympatry in Carangoidei. This study highlights the prevalence of secondary sympatry at a larger taxonomic scale than has previously been described in marine fishes. The prevalence of sympatry we observe coincides with evidence of morphological and environmental niche-partitioning in body size and depth in the water column between sister taxa. Our sister species biogeographic comparisons and trait analyses can only be completed with high taxonomic coverage and reliable estimates of phylogenetic relationships and species ranges. Fortunately, genomic sequencing techniques such as UCEs, combined with public databases such as IUCN and Aquamaps, allow us to address these critical questions across multiple taxa of marine organisms. Additional studies examining the mechanistic processes behind speciation in Carangoidei, including mate selection, reproductive timing, and mechanisms of dispersal at the species-level will shed further light on the drivers of speciation and biodiversity in this unique clade of marine fishes.

## Acknowledgements

We thank Gregory Watkins-Colwell of the Yale Peabody Museum of Natural History and Roger Bills of the South African Institute for Aquatic Biodiversity for their support. This research was supported by the Yale Institute for Biospheric Studies, the National Science Foundation Doctoral Dissertation Improvement Grant (DEB 1701597), the National Science Foundation Graduate Research Fellowship Program (DGE 1122492), the US Agency for International Development Research and Innovation Fellowship, the Yale Department of Ecology and Evolutionary Biology Chair’s Fund, and the Yale MacMillan Center International Dissertation Fellowship. PFC was also funded by the Australian Research Council (DE170100516). We are grateful to the following museum collections for providing samples for this study: Yale Peabody Museum of Natural History; University of Florida Board of Trustees – Florida Museum of Natural History – Genetic Resources Repository; South African Institute for Aquatic Biodiversity; Museums and Art Galleries of the Northern Territory, Fish Collection; University of Kansas Biodiversity Institute; Queensland Museum; Fisheries Research Laboratory, Mie University; Australian Museum; Academy of Natural Sciences of Drexel University; Louisiana State University Museum of Natural Science; Scripps Institution of Oceanography; and Kagoshima University Museum.

