## Supplemental_Information for "Widespread sympatry in a species-rich clade of marine fishes (Carangoidei)"

Supporting Information

**Methods**

Library amplification

We quantified libraries using a Qubit fluorometer after adapter ligation and cleanup, and we used 15µL of input library volume for amplification. The reaction mix consisted of 25 µL HiFi HotStart ReadyMix (KAPA Biosystems), 5 µL of indexed i5 and i7 primers (5 µM each), and 5 µL ddH20, with the following thermal profile: 98 °C for 45s; 12–14 cycles of 98 °C for 15s, 60 °C for 30s, 72 °C for 60s; and a final extension of 72 °C for 5m. The number of cycles was based on library concentrations measured after adapter ligation, as recommended by the KAPA Hyper Prep Library Kit protocol (KAPA Biosystems). We purified amplified libraries with a bead cleanup, quantified libraries using the Qubit fluorometer, and re-suspended clean libraries in 33 µL ddH20. We then pooled libraries into 11 groups of 8 or 9 libraries per pool at equimolar ratios, with a target concentration of 147 ng/µL (500 ng total DNA) for each of the 11 pools.

Hybridization

We ran the hybridization reaction at 65°C for 24 hours. Following hybridization, all pools were bound to streptavidin beads (MyOne C1; Life Technologies) and washed according to a standard target enrichment protocol (Blumenstiel et al. 2010). Samples were re-suspended with 30 µL ddH20 while still bound to the streptavidin beads, and we proceeded immediately to PCR recovery of the enriched libraries. The PCR protocol used 15 µL of streptavidin bead-bound, enriched library in water with 25 µL HiFi HotStart Taq (Kapa Biosystems), 5 µL of Illumina TruSeq primer mix (5 µM each), and 5 µL of ddH2O. We amplified each library using the following thermal profile: 98 °C for 45s; 16 cycles of 98 °C for 15s, 60 °C for 30s, 72 °C for 60s; and a final extension of 72 °C for 5m. We performed a 1X cleanup of the enriched DNA using Speed Beads and re-hydrated the resulting enriched DNA in 33 µL of ddH2O. To remove adapter-dimers <150 bp, we performed a clean-up step using the GeneRead Size selection kit (Qiagen, Inc.) on all enriched libraries, followed by quantification with a Qubit fluorometer. We estimated the size of the enriched pools by running each on an Agilent Bioanalyzer.

*phyluce* pipeline

Adapter sequences were trimmed from raw read data using the parallel wrapper (illumiprocessor) to implement the package *trimmomatic* (Bolger et al. 2014). We then assembled cleaned read data into contigs using *Trinity* vr2013-02-25 (Grabherr et al. 2013). To compute coverage across assemblies and summarize data by contig, we used a *phyluce* program (get_trinity_coverage.py) to 1) re-align the trimmed sequence reads to each set of assembled contigs using *bwa-mem* (Li 2013), 2) clean the resulting files and add read-group information using *picard* (1.99; http://picard.sourceforge.net/), 3) index the resulting files using *samtools* (Li et al. 2009), and 4) calculate coverage at each base of each assembled contig using *GATK* v2.7.2 (McKenna et al. 2010; DePristo et al. 2011; Van der Auwera et al. 2013). To identify assembled contigs that equate to enriched UCE loci, we used a *phyluce* program (match_contigs_to_loci.py) to align species-specific contig assemblies to a FASTA file containing the enrichment baits. The program removed duplicate contigs and contigs with baits targeting more than one UCE locus. It also filtered out matches less than 80% identical over 80% of the length of the contig. Subsequently, by inputting a file with the names of our taxa (Table S1), another *phyluce* program (get_match_counts.py) generated a list of UCE loci shared among all taxa ("complete" data matrix), as well as a list of UCE loci having data for any taxon ("incomplete" data matrix). We then used a program (get_fastas_from_match_counts.py) to generate separate monolithic FASTA files for the locus lists for the complete and incomplete data matrices. Next, we exploded each FASTA file (explode_get_fastas_file.py) by UCE locus, aligned and performed edge trimming on locus-specific FASTA files using MAFFT v7.130 (phyluce_align_seqcap_align.py; Katoh and Standley 2013) and converted the resulting files to NEXUS format.

**Table S1.** List of species and catalog information (institutional abbreviation and tissue number) used in this study. * indicates geographic range data were not included or not available. (1) indicates outgroup to Acanthomorpha. (2) indicates outgroup to Carangiformes. Catalog abbreviations are as follows: YFTC (Yale Fish Tissue Collection); UF (University of Florida Museum of Natural History, Fish Collection); SAIAB (South African Institute for Aquatic Biodiversity); MAGNT<AUS>:S (Museums and Art Galleries of the Northern Territory, Fish Collection); KUIT (University of Kansas Biodiversity Institute, Ichthyology Tissue Collection); QM (Queensland Museum); FRLM (Fisheries Research Laboratory Mie University); AM (Australian Museum); ANSP (Academy of Natural Sciences of Drexel University); LSUMZ (Louisiana State University, Museum of Natural Science); SIO (Scripps Institution of Oceanography); KAUM:I (Kagoshima University Museum, Ichthyology Collection).

| Species | Catalog Information | Species | Catalog Information |
|---|---|---|---|
| *Acanthurus bahianus*2* | YFTC 22882 | *Lampris guttatus*2* | YFTC 18113 |
| *Alectis alexandrina* | UF 236343 | *Lates calcarifer** | MAGNT<AUS>:S 92 |
| *Alectis ciliaris* | SAIAB ACEP09_742 | *Leptobrama muelleri** | UF 119722 |
| *Alectis indica* | SAIAB LS07-1029 | *Lichia amia* | SAIAB 127 |
| *Alepes apercna* | MAGNT<AUS>:S A01253 | *Makaira nigricans** | SIO 14_5 |
| *Alepes djedaba* | SAIAB AV2010-123 | *Megalaspis cordyla* | KUIT 4705 |
| *Alepes kleinii* | KUIT 8986 | *Mene maculata** | W.L. Smith 370; Pers. Coll. |
| *Alepes melanoptera* | UF 187084 | *Myripristis violacea*2* | YFTC 12630 |
| *Alepes vari* | QM I39514 | *Naucrates ductor* | SAIAB TZW-241 |
| *Apogon lateralis*2* | YFTC 12659 | *Nematistius pectoralis** | LSUMZ 7542 |
| *Atropus atropos* | LSUMZ 638 | *Oligoplites altus* | SIO 11_372 |
| *Atule mate* | SAIAB GG13_A011 | *Oligoplites saurus* | KUIT 10353 |
| *Carangichthyes dinema* | FRLM 40805 | *Pantolabus radiatus* | AM I34345_011 |
| *Carangoides armatus* | LSUMZ 13366 | *Parastromateus niger* | SAIAB SMITH 210.36_1 |
| *Carangoides bajad* | YFTC 40432 | *Phtheirichthys lineatus* | SAIAB ACEP09-224 |
| *Carangoides chrysophrys* | YFTC 40221 | *Polydactylus sexfilis** | KUIT 6829 |
| *Carangoides coeruleopinnatus* | SAIAB HMO7_005 | *Psettodes erumei** | LSUMZ 5295 |
| *Carangoides equula* | SAIAB ACEP08_1698 | *Pseudocaranx dentex* | SAIAB GG13_A024 |
| *Carangoides ferdau* | YFTC 40329 | *Pseudupeneus maculatus*2* | YFTC 19662 |
| *Carangoides fulvoguttatus* | SAIAB ACEP09_805 | *Rachycentron canadum* | SAIAB AV2010-184 |
| *Carangoides gymnostethus* | SAIAB JRG15_351 | *Remora albescens* | SIO 05_37 |
| *Carangoides hedlandensis* | MAGNT<AUS>:S OLD170 | *Remora brachyptera* | SIO 07_91 |
| *Carangoides humerosus* | AM I44858_022 | *Remora osteochir* | SIO 11_34 |
| *Carangoides malabaricus* | SAIAB LS07_0973 | *Remora remora* | SAIAB ACEP08-1660 |
| *Carangoides oblongus* | SAIAB ACEP08_1610 | *Scomber scombrus*2* | YFTC 13855 |
| *Carangoides orthogrammus* | SIO 11_81C | *Scomberoides commersonnianus* | SAIAB LS07-0891 |
| *Carangoides otrynter* | SIO 09_212 | *Scomberoides lysan* | LSUMZ 846 |
| *Carangoides plagiotaenia* | SAIAB T386 | *Scomberoides tala* | KAUM:I 59744 |
| *Carangoides praeustus* | FRLM 51405 | *Scomberoides tol* | SAIAB AV2010-091 |
| *Carangoides talamparoides* | LSUMZ 14022 | *Selar crumenophthalmus* | SAIAB HM07-317 |

| | | | |
|---|---|---|---|
| *Caranx bartholomaei* | ANSP 191545 | *Selaroides leptolepis* | KUIT 8987 |
| *Caranx bucculentus* | MAGNT<AUS>:S A00197 | *Selene brevoortii* | KUIT 8505 |
| *Caranx caballus* | SIO 14-6 | *Selene brownii* | KUIT 5855 |
| *Caranx caninus* | KUIT 8480 | *Selene dorsalis* | UF 236342 |
| *Caranx crysos* | YFTC 22281 | *Selene peruviana* | SIO 07_88L |
| *Caranx fischeri* | UF 236340 | *Selene setapinnis* | YFTC 13645 |
| *Caranx heberi* | YFTC 40413 | *Selene vomer* | YFTC 962 |
| *Caranx hippos* | YFTC 22225 | *Seriola dumerili* | SAIAB ACEP09_314 |
| *Caranx ignobilis* | YFTC 40404 | *Seriola fasciata* | KUIT 3260 |
| *Caranx latus* | UF 180908 | *Seriola hippos* | AM I44578_009 |
| *Caranx lugubris* | SAIAB PANGAEA-047 | *Seriola lalandi* | AM I46478_001 |
| *Caranx melampygus* | YFTC 28147 | *Seriola quinqueradiata* | KUIT 10317 |
| *Caranx papuensis* | YFTC 40216 | *Seriola rivoliana* | YFTC 22280 |
| *Caranx rhonchus* | SAIAB JRG15_292 | *Seriola zonata* | KUIT 1188 |
| *Caranx ruber* | YFTC 22276 | *Seriolina nigrofasciata* | SAIAB LS07-0815 |
| *Caranx senegallus* | UF 236341 | *Sphyraena putnamae** | KUIT 6785 |
| *Caranx sexfasciatus* | YFTC 40420 | *Sphyraena sphyraena** | UF LS499 |
| *Caranx tille* | YFTC 28124 | *Tetrapturus angustirostris** | SIO 05-31 |
| *Caranx vinctus* | LSUMZ 14570 | *Toxotes blythii** | YFTC 20205 |
| *Centropomus medius** | KUIT 8498 | *Toxotes jaculatrix** | LSUMZ 5166 |
| *Ceratoscopelus warmingii*1* | SIO 0510-09-28 | *Trachinotus africanus* | SAIAB AV2010-119 |
| *Chloroscombrus chrysurus* | YFTC 22286 | *Trachinotus anak* | QM I.38228 |
| *Chloroscombrus orqueta* | KUIT 8495 | *Trachinotus baillonii* | SAIAB RB09-160 |
| *Citharoides macrolepis** | KUIT 2468 | *Trachinotus blochii* | SAIAB TZW740 |
| *Coryphaena equiselis* | SAIAB ACEP08-1664 | *Trachinotus botla* | SAIAB AV2010-134 |
| *Coryphaena hippurus* | YFTC 40415 | *Trachinotus carolinus* | YFTC 11458 |
| *Decapterus akaadsi** | KAUM:I 75738 | *Trachinotus coppingeri* | AM I31254_004 |
| *Decapterus kurroides* | SAIAB HM07_034 | *Trachinotus goodei* | YFTC 1994 |
| *Decapterus macarellus* | SAIAB SMITH 210_26_1 | *Trachinotus kennedyi* | SIO 12_38 |
| *Decapterus macrosoma* | SAIAB HM07_346 | *Trachinotus mookalee* | UF 187093 |
| *Decapterus maruadsi* | KUIT 8984 | *Trachinotus paitensis* | SIO 07_91L |
| *Decapterus muroadsi* | SIO 11-377 | *Trachinotus rhodopus* | SIO 1181 |
| *Decapterus punctatus* | KUIT 1170 | *Trachurus capensis* | SAIAB F12 |
| *Decapterus russelli* | SAIAB HM07_071 | *Trachurus delagoa* | SAIAB AV2010-040 |
| *Decapterus smithvanizi** | KAUM:I 80712 | *Trachurus indicus** | AM I33820_040 |
| *Decapterus tabl* | SAIAB HM07-066 | *Trachurus japonicus* | KUIT 8597 |
| *Echeneis naucrates* | SAIAB HM07-581 | *Trachurus lathami* | YFTC 40427 |
| *Echeneis neucratoides* | KUIT 41346 | *Trachurus novaezelandiae* | AM I44627_012 |
| *Elagatis bipinnulata* | SAIAB T107 | *Trachurus symmetricus* | SIO 09_158 |
| *Gadus morhua*2* | YFTC 13930 | *Trachurus trachurus* | SAIAB JRG15-298 |
| *Gnathanodon speciosus* | SAIAB TZW-638 | *Trachurus trecae* | SAIAB JRG15-297 |

| | | | |
|---|---|---|---|
| *Hemicaranx amblyrhynchus* | KUIT 5142 | *Ulua aurochs* | QM I.38270 |
| *Hemicaranx zelotes* | SIO 09_269 | *Ulua mentalis* | YFTC 40155 |
| *Istiophorus platypterus** | KUIT 5428 | *Uraspis helvola* | SIO 10-20 |
| *Kajikia albida** | KUIT 5391 | *Uraspis secunda* | AM I46017 |
| *Kurtus gulliveri*2* | YFTC 18514 | *Uraspis uraspis* | LSUMZ 13829 |
| *Lactarius lactarius** | YFTC 25759 | *Xiphias gladius** | UF LS787 |

**Table S2.** Sample-size corrected Akaike information criterion (AICc) values used to compare three models of evolution: Brownian motion, Ornstein-Uhlenbeck and Early-burst.

| Trait | Brownian motion | Ornstein-Uhlenbeck | Early-burst |
|---|---|---|---|
| ln(maximum length) | 47.36806 | -16.63713 | 49.47023 |
| ln(maximum water column depth) | 227.9914 | 155.383 | 230.0936 |
| range size | 3050.682 | 3036.507 | 3052.784 |

**Table S3.** Number of sister species pairs consisting of two non-reef associated species (NR-NR), two reef associated species (R-R), and one reef-associated and one non-reef associated species (R-NR) grouped by A) all pairs, B) sympatric pairs and C) allopatric pairs.

| A) All Sister Species Pairs | |
|---|---|
| NR-NR | 14 |
| R-R | 14 |
| R-NR | 13 |
| **B) Sympatric Sister Species Pairs** | |
| NR-NR | 6 |
| R-R | 12 |
| R-NR | 12 |
| **C) Allopatric Sister Species Pairs** | |
| NR-NR | 8 |
| R-R | 2 |
| R-NR | 1 |

**Table S4.** Sources for fish images from Figure 2. Species are listed in order they appear in Figure 2, from top to bottom. CSIRO is the abbreviation for Australian National Fish Collection (CC BY 3.0 AU). All images corresponding to names were obtained with permission from FishBase (CC BY-NC 3.0).

| Species | Image Source |
|---|---|
| *Seriola dumerili* | JE Randall |
| *Seriola rivoliana* | CSIRO |
| *Seriola lalandi* | CSIRO |
| *Trachinotus anak* | CSIRO |
| *Trachinotus blochii* | JE Randall |
| *Trachinotus baillonii* | CSIRO |
| *Scomberoides tala* | CSIRO |
| *Scomberoides commersonnianus* | CSIRO |
| *Remora brachyptera* | CSIRO |
| *Phtheirichthys lineatus* | CSIRO |
| *Echeneis naucrates* | JE Randall |
| *Coryphaena* sp. | JE Randall |
| *Istiophorous platypterus* | CSIRO |
| *Mene maculatus* | JE Randall |
| *Leptobrama muelleri* | JE Randall |
| *Sphyraena* sp. | JE Randall |
| *Lactarius lactarius* | CSIRO |
| *Centropomus medius* | JE Randall |
| *Caranx papuensis* | CSIRO |
| *Caranx ignobilis* | R Daly |
| *Megalaspis cordyla* | CSIRO |
| *Caranx bucculentus* | CSIRO |
| *Caranx ruber* | TA Meyer |
| *Pantalobus radiatus* | CSIRO |
| *Alepes vari* | CSIRO |
| *Chloroscombrus chrysurus* | C Meiners |
| *Selaroides leptolepis* | CSIRO |
| *Carangoides fulvoguttatus* | CSIRO |
| *Carangoides coeruleopinnatus* | CSIRO |
| *Ulua aurochs* | CSIRO |
| *Carangoides hedlandensis* | CSIRO |
| *Selene vomer* | T Meyer |
| *Alectis indica* | JE Randall |
| *Uraspis uraspis* | CSIRO |
| *Carangoides ferdau* | CSIRO |
| *Decapterus macrosoma* | CSIRO |
| *Decapterus muroadsi* | CSIRO |
| *Decapterus macarellus* | JE Randall |
| *Decapterus akaadsi* | JE Randall |

| | |
|---|---|
| *Trachurus indicus* | JE Randall |
| *Trachurus trachurus* | M Garc |
| *Carangoides equula* | CSIRO |

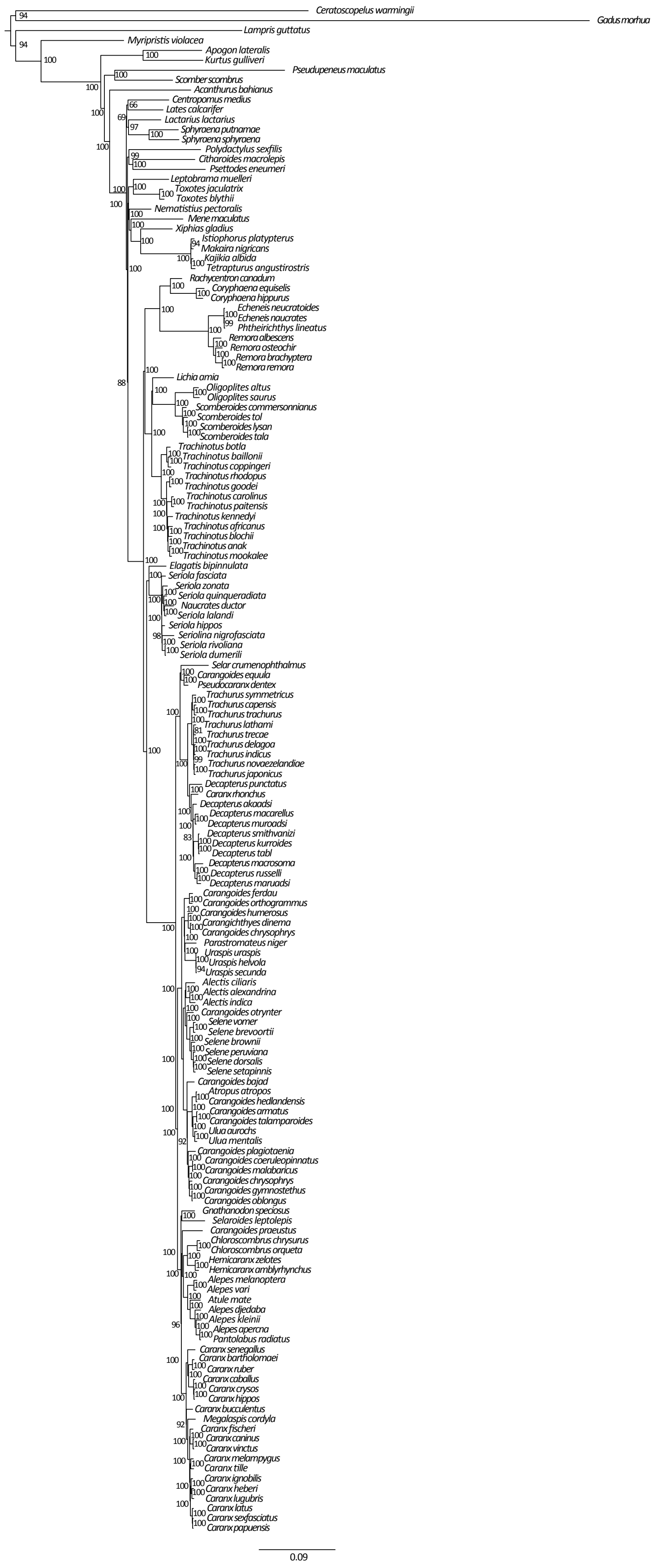


**Figure S1**. Maximum likelihood phylogeny of 154 Carangiformes and outgroup taxa constructed in IQ-TREE using a 75% complete matrix. Bootstrap support values are indicated for each node.

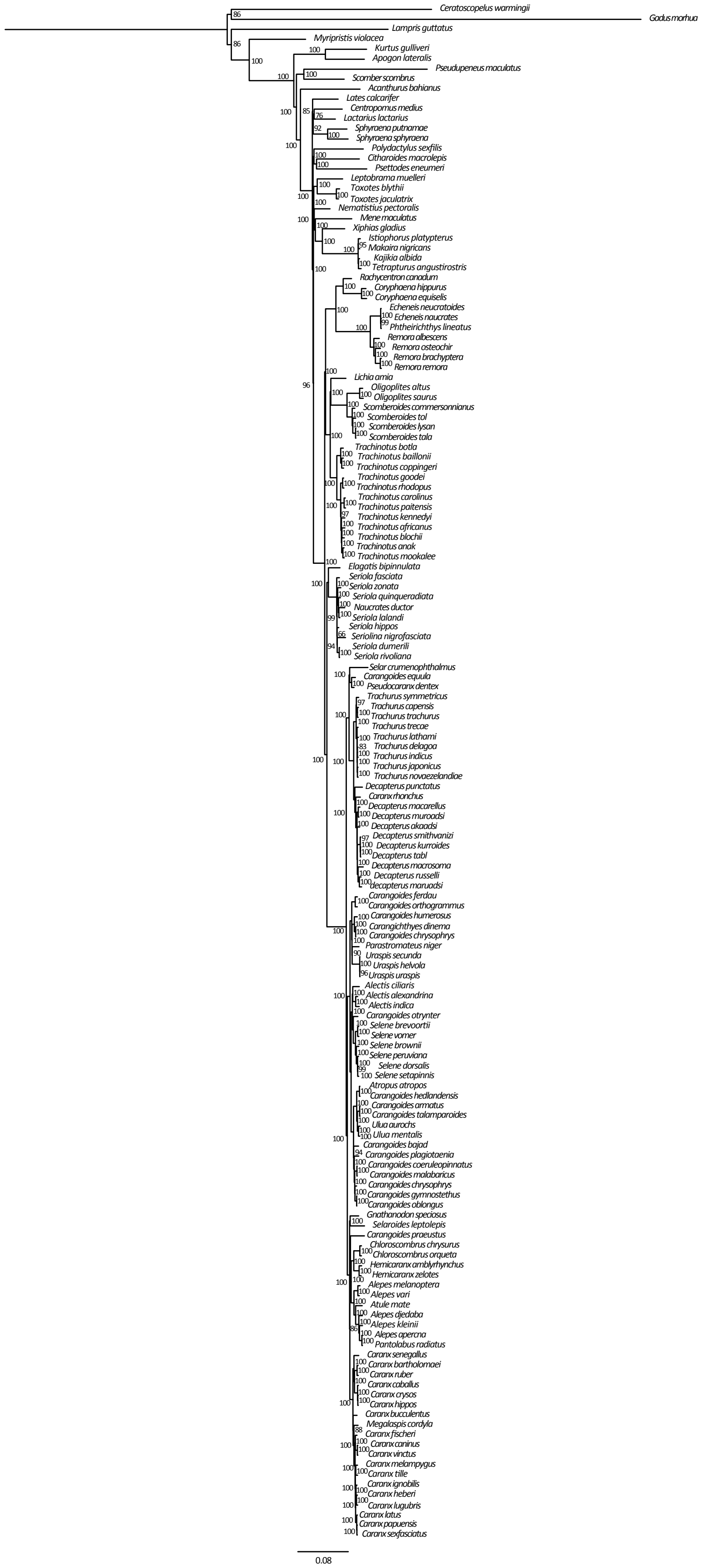

**Figure S2**. Maximum likelihood phylogeny of 154 Carangiformes and outgroup taxa constructed in IQ-TREE using a 95% complete matrix. Bootstrap support values are indicated for each node.

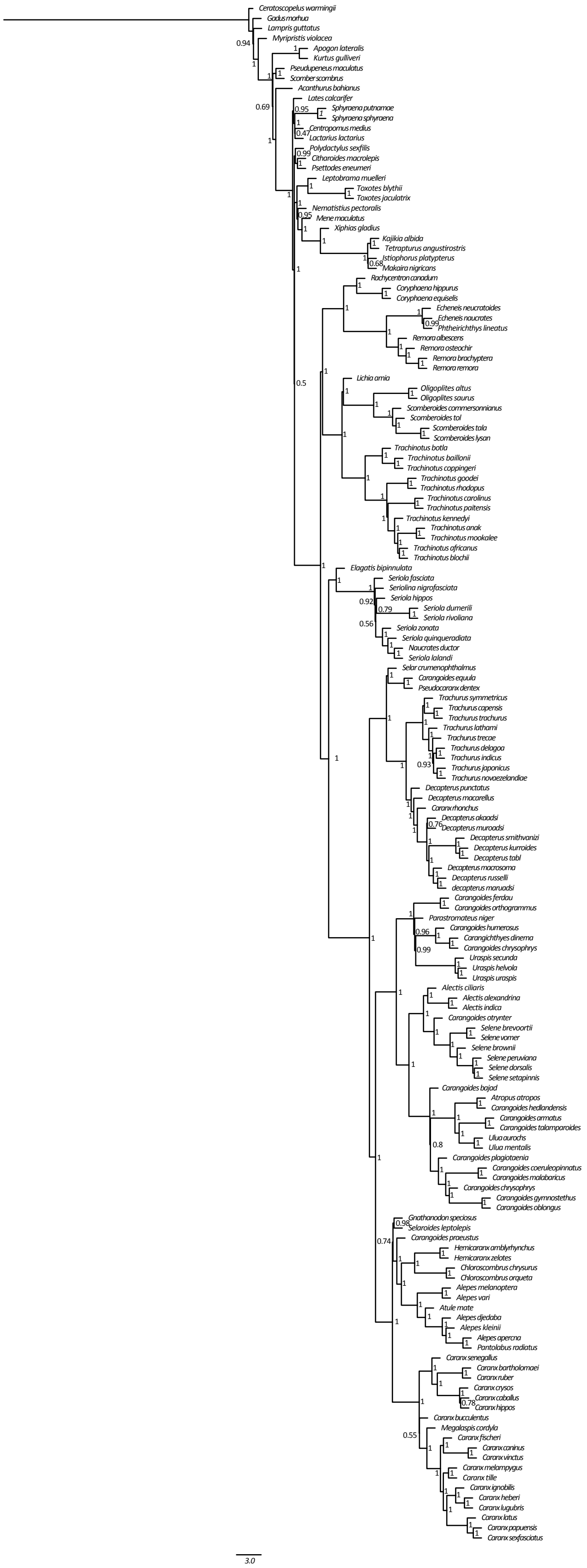

**Figure S3**. Majority rule consensus tree of 154 Carangiformes and outgroup taxa generated in ASTRAL-II using loci from the 75% complete matrix after generating individual gene trees in MrBayes v3.2.6. ASTRAL support values are indicated for each node.

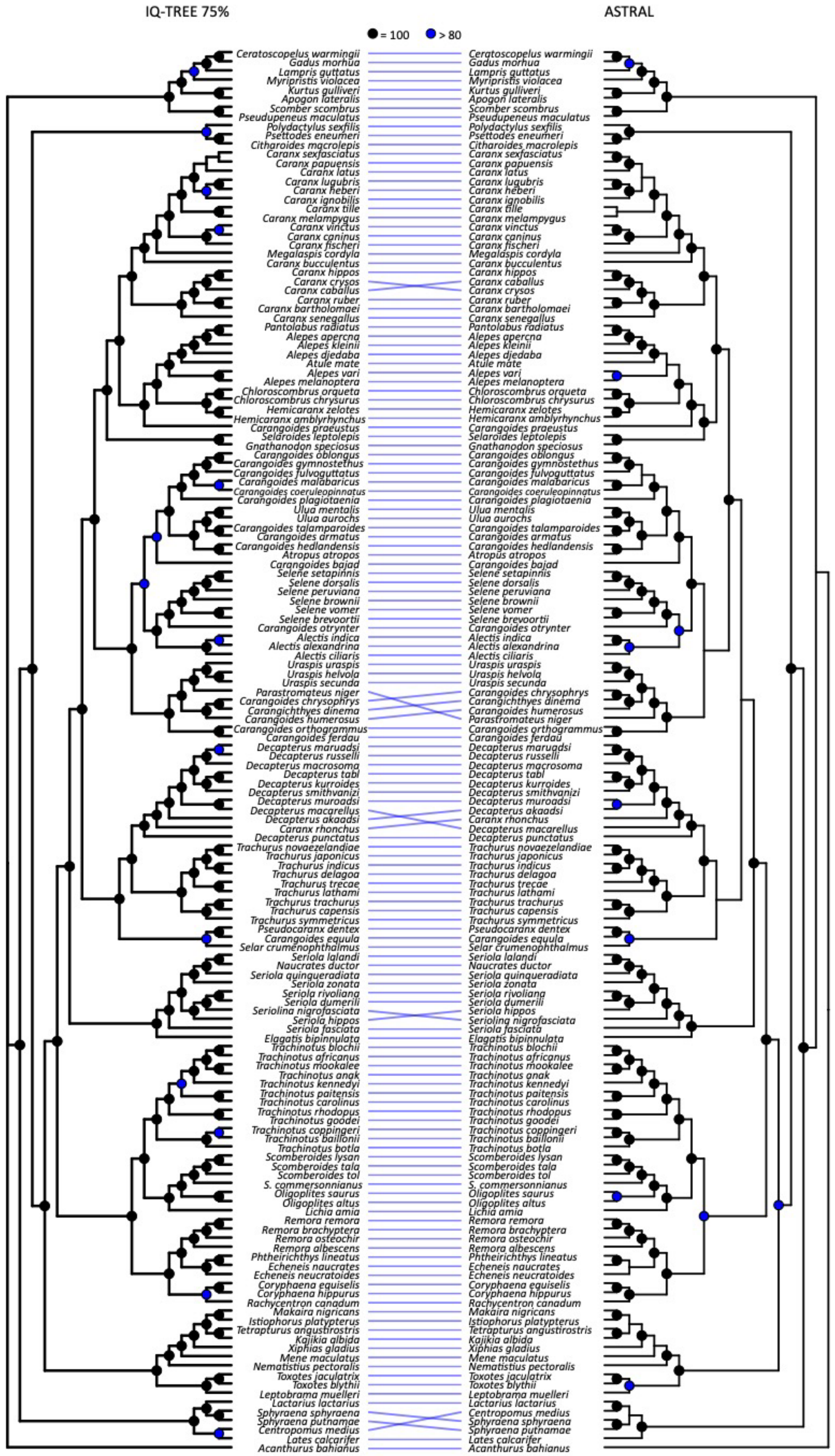


**Figure S4**. Tanglegram of maximum likelihood phylogeny constructed in IQ-TREE using a 75% complete matrix compared to a majority rule consensus tree generated in ASTRAL-II.

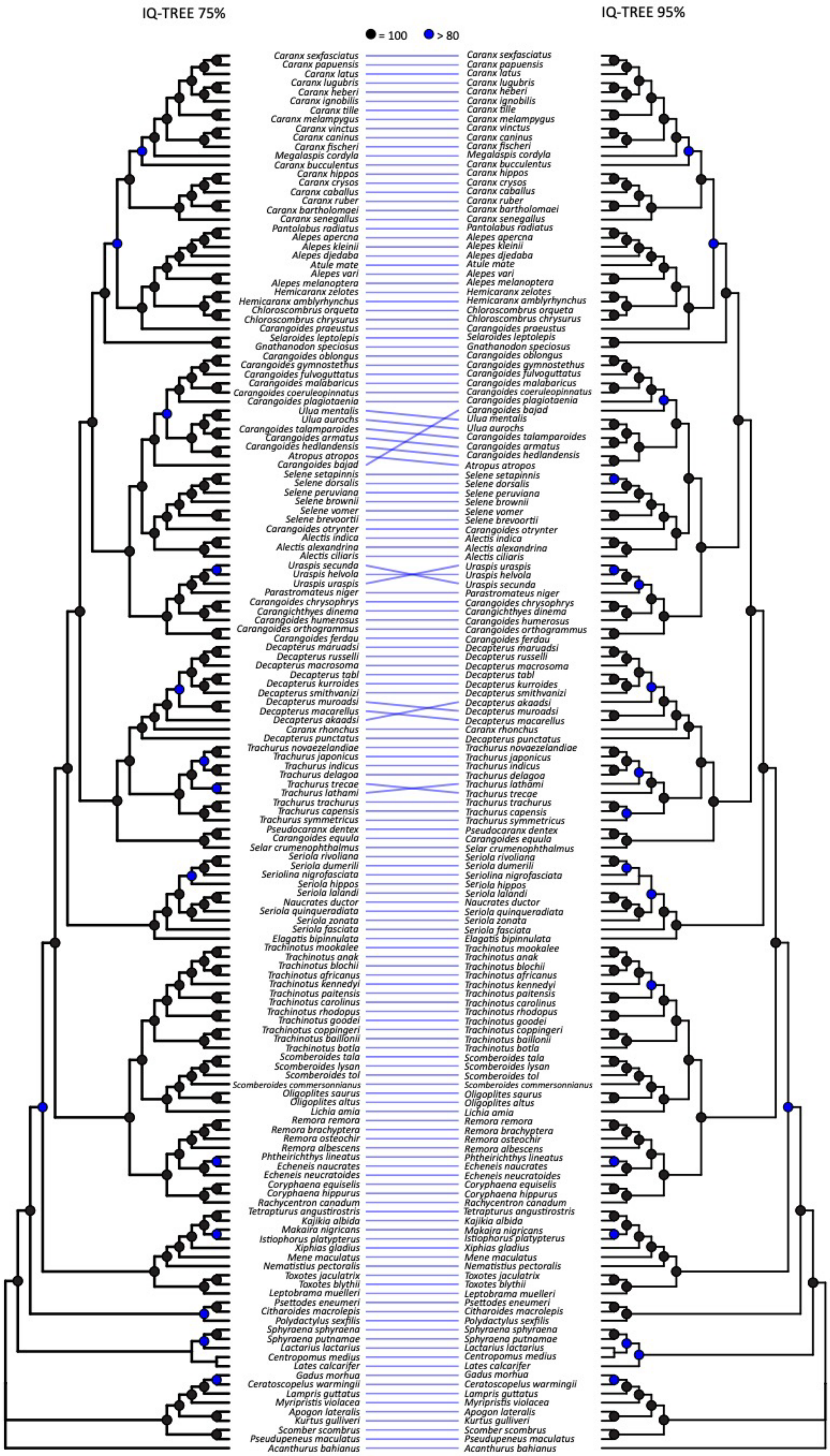


**Figure S5**. Tanglegram of maximum likelihood phylogenies constructed in IQ-TREE using a 75% complete matrix and a 95% complete matrix.

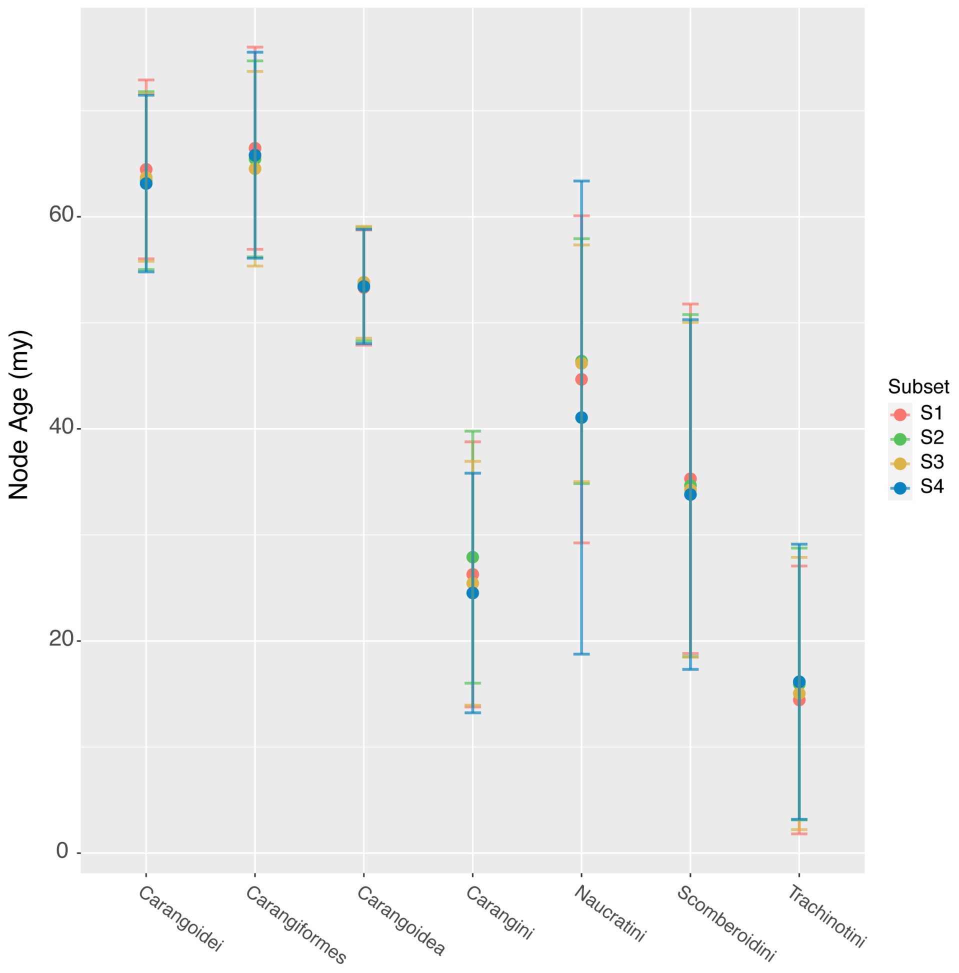


**Figure S6.** Divergence date estimates and 95% High Posterior Densities for selected Carangiformes nodes, generated in BEAST v1.10.4. We ran four subsets using random samples of 25 UCE loci.

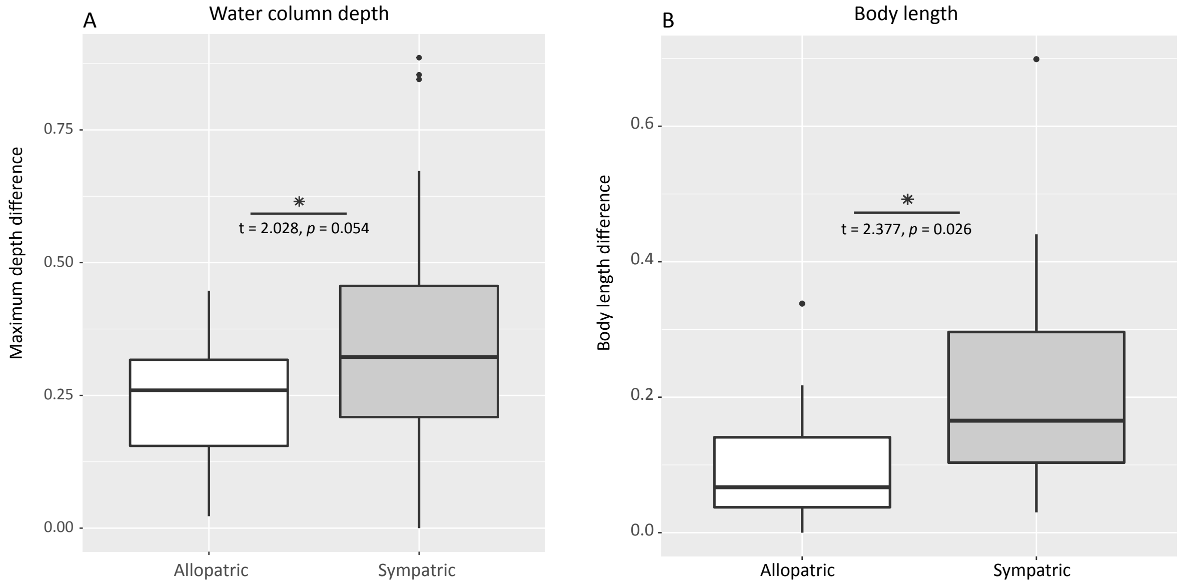


**Figure S7.** Contrasts between allopatric and sympatric sister species pairs for maximum water column depth (A) and maximum body length (B) for 37 sister species pairs after removing four pairs due to conflicting tree topologies between the phylogeny constructed with the IQ-TREE 75% complete matrix versus the ASTRAL majority rule consensus tree. Results of Welch's *t*-tests are presented to show significance between allopatric and sympatric sister species pairs.

### Fossil Calibrations

We included the following nine fossil calibration points as described in Harrington, R. C., B. C. Faircloth, R. I. Eytan, W. L. Smith, T. J. Near, M. E. Alfaro, and M. A. Friedman. 2016. Phylogenomic analysis of carangimorph fishes reveals flatfish asymmetry arose in a blink of the evolutionary eye. BMC Evol. Biol. 16:224.

#### Calibration 1

**Node calibrated.** MRCA of *Lampris guttatus* and *Myripristis violacea*.

**Fossil taxon and specimen.** *Aipichthys minor*, MNHN HDJ 65, Museum national d'Histoire Naturelle, Paris.

**Minimum age.** 98.0 Ma.

**Phylogenetic justification.** *Aipichthys minor* is placed on the lampridiform stem by Davesne et al. (2014) in a parsimony analysis of 67 morphological characters.

**Age justification.** Age evidence for the fish beds at Hadjula is reviewed elsewhere (Benton et al. 2015), but key details are provided for completeness. This horizon is located below reported occurrence of *Mantelliceras mantelli*, which defines the first complete ammonite zone of the Late Cretaceous. The top of the *Mantelliceras mantelli* Zone is dated at 98.0 Ma (Ogg et al. 2012), from which we derive a minimum age for *Aipichthys*.

**Outgroup age sequence.** 247.1, 236.0, 221.0, 193.81, 181.7, 166.1, 151.2, 150.94, 150.94, 125, 98.0

**Estimated constraints on node age prior.** Mean: 119.1 Ma; 95% CI: 143.0 Ma.

#### Calibration 2

**Node calibrated.** MRCA of *Myripristis violacea* and *Kurtus gulliveri*.

**Fossil taxon and specimen.** *Stichocentrus liratus*, NHMUK PV P.47835 (holotype), The Natural History Museum, London, UK.

**Minimum age.** 98.0 Ma.

**Phylogenetic justification.** The penultimate anal-fin spine of *Stichocentrus* is enlarged, and represents a synapomorphy of holocentroids (Zehren 1979; Patterson 1993).

**Age justification.** Age evidence for the fish beds at Hadjula is reviewed elsewhere (Benton et al. 2015), but key details are provided for completeness. This horizon is located below reported occurrence of *Mantelliceras mantelli*, which defines the first complete ammonite zone of the Late Cretaceous. The top of the *Mantelliceras mantelli* Zone is dated at 98.0 Ma (Ogg et al. 2012), from which we derive a minimum age for *Stichocentrus*.

**Outgroup age sequence.** 247.1, 236.0, 221.0, 193.81, 181.7, 166.1, 151.2, 150.94, 150.94, 125, 98.0, 98.0

**Estimated constraints on node age prior.** Mean: 108.7 Ma; 95% CI: 128.8 Ma.

**Calibration 3**

**Node calibrated.** MRCA of *Lates calcarifer* and *Centropomus medius*.

**Fossil taxon and specimen.** *Eolates gracilis*, MNHN BOL 61, 62 (holotype), Museum national d'Histoire Naturelle, Paris.

**Minimum age.** 49 Ma.

**Phylogenetic justification.** *Eolates gracilis* is resolved as a sister lineage of *Lates* to the exclusion of *Centropomus* in a parsimony analysis of 29 morphological characters (Otero 2004).

**Age justification.** *Eolates gracilis* is known from the Pesciara locality of Bolca, Italy. A detailed review of the geology and age of this deposit is given by Papazzoni and colleagues (Papazzoni et al. 2014), but key details are summarized here. Pesciara can be constrained to the narrow interval of overlap between NP14 and SBZ11. This constrains the deposits to no younger than 49 Ma (Vandenberghe et al. 2012), which we apply as a minimum age here.

**Outgroup age sequence.** 247.1, 236.0, 221.0, 193.81, 181.7, 166.1, 151.2, 150.94, 150.94, 125, 98.0, 98.0, 69.71, 69.71, 55.2, 49

**Estimated constraints on node age prior.** Mean: 58.3 Ma; 95% CI: 72.8 Ma.

**Calibration 4**

**Node calibrated.** MRCA of *Mene maculata* and *Xiphias gladius*.

**Fossil taxon and specimen.** *Mene purydi*, USNM 494403 (holotype and only specimen), National Museum of Natural History, Washington, DC, USA.

**Minimum age.** 55.20 Ma.

**Phylogenetic justification.** Friedman and Johnson (2005) interpret *Mene purdyi* as a menid on the based on a mixture of derived and general traits. *M. purdyi* shares three compelling synapomorphies with *Mene*: a cavernous vault formed by the frontals; an infraorbital series comprising numerous, small ossicles; and close application of the neural arches of the first two vertebrae.

**Outgroup age sequence.** Friedman and Johnson (2005) identified *Mene purdyi* as latest Thanetian-earliest Ypresian in age on the basis of a series of plantonic foraminifera collected from the matrix of the specimen consistent with foraminiferal zones P4c to P5. Specifically, the presence of *Morozovella velascoensis* provides an upper age limit of 55.20 Ma (Anthonissen and Ogg, 2012).

**Calibration prior.** 247.1, 236.0, 221.0, 193.81, 181.7, 166.1, 151.2, 150.94, 150.94, 125, 98.0, 98.0, 69.71, 69.71, 55.2

**Estimated constraints on node age prior.** Mean: 67.5 Ma; 95% CI: 84.7 Ma.

**Notes.** Roughly contemporary specimens of *Mene* are known from the Stolleklint Clay of Denmark (Bonde et al. 2008) and the Danatinsk Formation of Turkmenistan (Bannikov 2010; Bannikov et al. 1997). Available evidence constrains the minimum age of both deposits to earliest Eocene, so we adopt *Mene purdyi* as our calibration here.

### Calibration 5

**Node calibrated.** MRCA of *Echeneis* cf. *naucratoides* and *Rachycentron canadum*.

**Fossil taxon and specimen.** Echeneidae undet., HLMD WT-36, Hessisches Landesmuseum, Darmstadt, Germany (Friedman et al. 2014).

**Minimum age.** 29.62 Ma.

**Phylogenetic justification.** Echeneidae undet. bears numerous synapomorphies of remoras, including a dorsal adhesion disc and expanded transverse processes of vertebrae (Friedman et al. 2013; O'Toole 2002).

**Age justification.** The 'fish shales' of Grube Unterfeld ("Frauenweiler") yielding Echenidae undet. lie within NP23 (Sakamoto et al. 2004). The top of NP23 is dated to 29.62 Ma (Vandenberghe et al. 2012), providing a minimum age of divergence between *Echeneis* cf. *naucratoides* and *Rachycentron canadum*.

**Outgroup age sequence.** 247.1, 236.0, 221.0, 193.81, 181.7, 166.1, 151.2, 150.94, 150.94, 125, 98.0, 98.0, 69.71, 69.71, 55.2, 54.17, 49.0, 49.0, 29.62

**Estimated constraints on node age prior.** Mean: 41.0 Ma; 95% CI: 51.9 Ma.

**Notes.** The Rupelian remora †*Opisthomyzon* bears an adhesion disc that is more primitive than those of Echeneidae undet. (Friedman et al. 2013). Although the maximum age of this genus is given by radiometric dating of underlying sediments, its minimum age is not as well constrained (Friedman et al. 2013); foraminiferans have not been reported from the deposits yielding †*Opisthomyzon* (Furrer and Leu 1998). Because the minimum age of Echeneidae undet. can be constrained through foraminiferal biostratigraphy (see above), we have selected this fossil as our calibration rather than the better known †*Opisthomyzon*.

**Calibration 6**

**Node calibrated.** MRCA of *Echeneis* cf. *naucratoides* and *Scomberoides commersonnianus*.

**Fossil taxon and specimen.** *Ductor vestenae*, MNHN BOL 96, Museum national d'Histoire Naturelle, Paris.

**Minimum age.** 49 Ma.

**Phylogenetic justification.** Based on a series of parsimony and Bayesian analyses of morphological plus molecular data and morphological data in isolation, Friedman et al. (2013) reported two alternative placements for *Ductor*: either as the sister taxon of crown Echeneoidei, or within crown Echeneoidei as sister to Rachycentridae plus Coryphaenidae. We adopt the former, interpretation here, as it represents a more conservative application of this fossil as a minimum.

**Age justification.** *Ductor vestenae* is known from the Pesciara locality of Bolca, Italy. A detailed review of the geology and age of this deposit is given by Papazzoni and colleagues (Papazzoni et al. 2014), but key details are summarized here. Pesciara can be constrained to the narrow interval of overlap between NP14 and SBZ11. This constrains the deposits to no younger than 49 Ma (Vandenberghe et al. 2012), which we apply as a minimum age here.

**Outgroup age sequence.** 247.1, 236.0, 221.0, 193.81, 181.7, 166.1, 151.2, 150.94, 150.94, 125, 98.0, 98.0, 69.71, 69.71, 55.2, 54.17, 49.0, 49.0

**Estimated constraints on node age prior.** Mean: 52.2 Ma; 95% CI: 59.1 Ma.

**Calibration 7**

**Node calibrated.** MRCA of *Scomberoides commersonnianus* and *Trachinotus blochii*.

**Fossil taxon and specimen.** *Scomberoides spinosus*, PIN 485/72, Paleontological Institute of the Russian Academy of Sciences, Moscow, Russia.

**Minimum age.** 19.30 Ma.

**Phylogenetic justification.** *Scomberoides spinosus* shows two key hard-tissue synapomorphies of Scomberoidini (Smith-Vaniz 1984; Gushiken 1988): 26 vertebrae (other carangids generally have 24), and posterior fin rays of the dorsal and anal fin developed as finlets (Bannikov 1990).

**Age justification.** *Scomberoides spinosus* derives from sediments of the Upper Maikop at Chernaya Rechka, Caucasus (Bannikov and Parin 1997). These deposits are placed within the Sakaraul regional stage, which is correlated with the upper part of Plantonic Foraminiferan Zone M2 (Rögl 1998). The top of M2 is dated as 19.30 Ma (Hilgen et al. 2012), which we adopt as a minimum age for the divergence between *Scomberoides commersonnianus* and *Trachnotus blochii*.

**Outgroup age sequence.** 247.1, 236.0, 221.0, 193.81, 181.7, 166.1, 151.2, 150.94, 150.94, 125, 98.0, 98.0, 69.71, 69.71, 55.2, 54.17, 49.0, 49.0, 19.30

**Estimated constraints on node age prior.** Mean: 35.8 Ma; 95% CI: 50.9 Ma.

**Notes.** The age estimate provided here is a minor adjustment from previous applications of this calibration. Bannikov (2010) has interpreted the Eocene (Bartonian) *Quasioligoplites mirus* as a member of Scomberoidini, which, if correct, means that our proposed minimum is a substantial underestimate of the divergence between *Scomberoides commersonnianus* and *Trachinotus blochii*. Although *Quasioligoplites mirus* broadly resembles a scomberoidine, it does not clearly show diagnostic hard-tissue characters of this clade that are readily apparent in *Scomberoides spinosus* (Bannikov 1990), which we regard as a more conservative marker.

**Calibration 8**

**Node calibrated.** MRCA of *Seriola zonata* and *Chloroscombrus orqueta*.

**Fossil taxon and specimen.** *Eastmanalepes primaevus*, MCZ 50706 (holotype), Museum of Comparative Zoology, Harvard University, Cambridge, USA.

**Minimum age.** 49 Ma.

**Phylogenetic justification.** *Eastmanelepes* bears thickened scutes along its flank, representing a synapomorphy of Carangini within Carangidae (Smith-Vaniz 1984; Gushiken 1988).

**Age justification.** *Eastmanelepes* is known from the Pesciara locality of Bolca, Italy. A detailed review of the geology and age of this deposit is given by Papazzoni and colleagues (2014), but key details are summarized here. Pesciara can be constrained to the narrow interval of overlap between NP14 and SBZ11. This constrains the deposits to no younger than 49 Ma (Vandenberghe et al. 2012), which we apply as a minimum age here.

**Outgroup age sequence.** 247.1, 236.0, 221.0, 193.81, 181.7, 166.1, 151.2, 150.94, 150.94, 125, 98.0, 98.0, 69.71, 69.71, 55.2, 54.17, 49.0, 49.0

**Estimated constraints on node age prior.** Mean: 52.2 Ma; 95% CI: 59.1 Ma.

**Calibration 9**

**Node calibrated.** MRCA of *Psettodes erumei* and *Polydactylus sexfilis*.

**Fossil taxon and specimen.** *Heteronectes chaneti*, NHMW 1974.1639.24, 1974.1639.25 (holotype and only specimen), Naturhistorisches Museum, Vienna, Austria.

**Minimum age.** 49 Ma.

**Phylogenetic justification.** Friedman (2008) resolved *Heteronectes* as the deepest branch of the flatfish stem lineage in a parsimony analysis of 58 morphological characters.

**Age justification.** *Heteronectes* is known from the fish beds of Bolca, Italy, but whether it derives from the Pesciara or Monte Postale locality is unclear. A detailed review of the geology and age of this deposit is given by Papazzoni and colleagues (Papazzoni et al. 2014), but key details are summarized here. Pesciara can be constrained to the narrow interval of overlap between NP14 and SBZ11. This constrains the deposits to no younger than 49 Ma (Vandenberghe et al. 2012), which we apply as a minimum age here.

**Outgroup age sequence.** 247.1, 236.0, 221.0, 193.81, 181.7, 166.1, 151.2, 150.94, 150.94, 125, 98.0, 98.0, 69.71, 69.71, 55.2, 49.0

**Estimated constraints on node age prior.** Mean: 58.3 Ma; 95% CI: 72.8 Ma.
